# Receptor-like kinase RLK15 binds a fungal RNase effector to enhance autophosphorylation and activate host defense

**DOI:** 10.64898/2026.10.15.759029

**Authors:** Lei Li, Yuhe Lian, Li Zhang, Huan Jia, Yunpeng Gai, Guy Smagghe, Song Tang, Mengmeng Yang, Enping Cai, Yizhen Deng, Zide Jiang, Lianhui Zhang, Xiaofan Zhou, Changqing Chang

## Abstract

Fungal pathogens secrete diverse effectors to suppress host immunity, yet the plant receptors that perceive these virulence factors often remain elusive. Here, we identify Ss1624, a secreted RNase from the sugarcane smut pathogen *Sporisorium scitamineum*, and show that it is recognized by the plasma-membrane-localized CrRLK1L-family receptor kinase RLK15. This recognition triggers canonical immune outputs, including reactive oxygen species burst, callose deposition, defense gene activation, and induces hypersensitive cell death. Mechanistically, Ss1624 physically interacts with the receptor-like kinase RLK15, which is required for Ss1624-triggered cell death. Functional assays indicate that this interaction enhances RLK15 autophosphorylation in a kinase activity-dependent manner, suggesting that Ss1624 can act as an elicitor. Transcript profiling further reveals coordinated expression patterns of Ss1624 and RLK15 during infection, and perturbation of Ss1624 expression alters RLK15 transcript levels and pathogenic outcomes. Extending these findings to rice, the ortholog OsRLK15 contributes positively to resistance to bacterial and fungal diseases without yield penalty in rice. Our results support a model in which a secreted RNase effector is directly sensed by a receptor kinase to initiate broad-spectrum immune responses, providing insights into host-pathogen interactions and identifying RLK15 as a candidate for crop improvement.

**Short Summary:** A secreted RNase from *Sporisorium scitamineum*, is recognized by receptor kinase RLK15, triggering cell death and immunity. Ss1624 binds RLK15 and promotes its autophosphorylation in a kinase-activity-dependent manner. Transcript analysis reveals coordinated expression of the effector and receptor during infection. Notably, the rice ortholog OsRLK15 confers broad-spectrum disease resistance without yield penalty. These findings identify RLK15 as a conserved immune receptor for RNase-type effectors and a promising target for engineering durable crop resistance.

## Introduction

Plants rely on layered immune surveillance to detect microbes while preserving the receptor signaling needed for growth and development.(Jian *et al*., 2024) This discrimination is especially challenging for secreted pathogen proteins, because molecules that contribute to colonization can also expose the pathogen to host recognition. Understanding how plants convert such pathogen-derived proteins into receptor activation is therefore central to explaining the evolutionary boundary between virulence and immunogenicity.

At the cell surface, pattern-recognition receptors (PRRs) perceive microbial or damage-associated ligands and initiate pattern-triggered immunity (PTI), whereas intracellular nucleotide-binding leucine-rich repeat receptors commonly detect pathogen effectors and activate effector-triggered immunity (ETI).(Chen *et al*., 2025) Canonical PRRs such as FLS2, EFR and PEPR signal through receptor complexes that include BAK1.(Gómez-Gómez & Boller, 2000; Chinchilla *et al*., 2006; Yamaguchi *et al*., 2006; Zipfel *et al*., 2006) These immune branches are now understood to reinforce one another rather than operate as isolated layers. A key unresolved question is how secreted effector proteins that are not canonical microbial patterns can themselves become immunogenic signals, particularly when recognition is coupled directly to receptor-kinase activation.

Cell-surface receptor kinases provide the first signaling interface for many extracellular immune cues, triggering outputs such as reactive oxygen species (ROS) production, callose deposition and transcriptional reprogramming.(Couto & Zipfel, 2016; Dodds *et al*., 2024; Ma, M *et al*., 2025) Pathogens counter this surveillance with secreted effectors that suppress or redirect host signaling (Djamei *et al*., 2011; Kleemann *et al*., 2012; Liu *et al*., 2025; Wei *et al*., 2025) Although BAK1- and SOBIR1-centered receptor complexes account for many characterized cell-surface immune responses,(Gómez-Gómez & Boller, 2000; Chinchilla *et al*., 2006; Zipfel *et al*., 2006; Yamaguchi *et al*., 2010) the diversity of receptor architectures implies that alternative routes of effector perception and kinase activation remain to be defined.

Smut fungi represent a major threat to global food and bioenergy security, causing devastating diseases in vital gramineous crops such as sugarcane, maize, and wheat.(Schulze-Lefert & Panstruga, 2003; Zuo *et al*., 2019) During infection, smut fungi secrete diverse effectors to manipulate host immunity.(Ma *et al*., 2018; Ökmen *et al*., 2022; Saado *et al*., 2022; Ling *et al*., 2025; Lee *et al*., 2026) While many effectors function as virulence factors, accumulating evidence suggests that certain effectors may also act as conditional avirulence determinants, depending on whether the plant possesses the corresponding receptor.(Praz *et al*., 2017) In the sugarcane smut fungus *Sporisorium scitamineum*, genomic analysis has identified 464 predicted secreted proteins, yet nearly half of which lack functional annotation.(Schuster *et al*., 2018) Elucidating the pathogenic roles of these proteins and their cognate host targets is therefore essential for deciphering the molecular trade-offs underlying smut disease.

Secreted ribonuclease (RNase)-type proteins illustrate this functional diversity. Some RNase effectors promote virulence; for instance, deletion of effectors such as Fg12 (*Fusarium graminearum*), FoRnt2 (*Fusarium oxysporum*), SRE1 (*Setosphaeria turcica*), Nuc1, and Nuc2 (*Ustilago maydis*) respectively reduces the fungal virulence, highlighting their pro-pathogenic function.(Mukherjee *et al*., 2020; Yang *et al*., 2021; He *et al*., 2022; Qian *et al*., 2022) Conversely, the removal of VdRTX1 (*Verticillium dahliae*), SRN1, and SRN2 (*Colletotrichum orbiculare*) enhances the fungal virulence, indicating that RNase proteins may also expose pathogens to host immunity.(Yin *et al*., 2022; Kumakura *et al*., 2024) Thus, the central mechanistic question is not simply whether fungal RNases affect immunity, but how a host receptor can connect perception of an immunogenic RNase-type effector to receptor activation and downstream defense.

Here we identify Ss1624, an RNase T1-like effector from *S. scitamineum*, as a pathogen protein that is both enzymatically active and immunogenic. Ss1624 induces multiple defense-associated outputs and its expression level is inversely associated with smut virulence. We identify the CrRLK1L-family receptor kinase RLK15 as an Ss1624-associated host protein and show genetically that RLK15 homologues are required for Ss1624-triggered cell death in *N. benthamiana.* Structure-function analyses further link RLK15 membrane localization and kinase competence to immune activity and detectable Ss1624 association, while purified Ss1624 enhances autophosphorylation of the RLK15 kinase domain *in vitro*. Moreover, modulation of Ss*1624* expression influences RLK15 induction and impacts virulence outcomes, underscoring the functional relevance of this interaction. Importantly, the orthologous receptor OsRLK15 in rice positively contributes to resistance against both bacterial and fungal pathogens, suggesting evolutionary conservation and potential translational relevance.

Collectively, our findings demonstrate that RNase effectors can also serve as immunogenic signals. By revealing receptor-mediated recognition of secreted RNases, this work not only expands the conceptual boundaries of PTI and ETI integration but also establishes RNase effectors themselves as pathogen-derived immunogenic features.

## Result

### Ss1624 is an immunogenic fungal RNase whose abundance restrains smut virulence

To identify functionally relevant effectors in *S. scitamineum*, we analyzed the *S. scitamineum* BSES genome project (PRJEB5169) (Taniguti *et al*., 2015) with SignalP- 5.0, TMHMM-2.0, and EffectorP-3.0, which yielded 98 putative canonical effectors with predicted N-terminal signal peptides (SPs) and no transmembrane domains. Integrating with RNA-seq datasets from infected sugarcane identified 47 candidates that were differentially expressed during infection. Transient expression in *N. benthamiana* identified one protein, designated Ss1624 (GenBank: CDW99538.1), that triggered cell death and contained a predicted N-terminal SP and an RNase T1-like domain (**Figure 1A**). A yeast signal sequence trap assay confirmed the functionality of its SP for secretion (**Supplemental Figures 1A and 1B**). Recombinant Ss1624 purified from *P. pastoris* degraded total RNA from both dicot and monocot plants *in vitro* (**Supplemental Figures 1C-1E**), establishing Ss1624 as a secreted fungal ribonuclease. When transiently expressed in *N. benthamiana*, Ss1624-RFP accumulated at the plasma-membrane region and in the nucleus (**Supplemental Figure 1F**).

**Figure 1.**
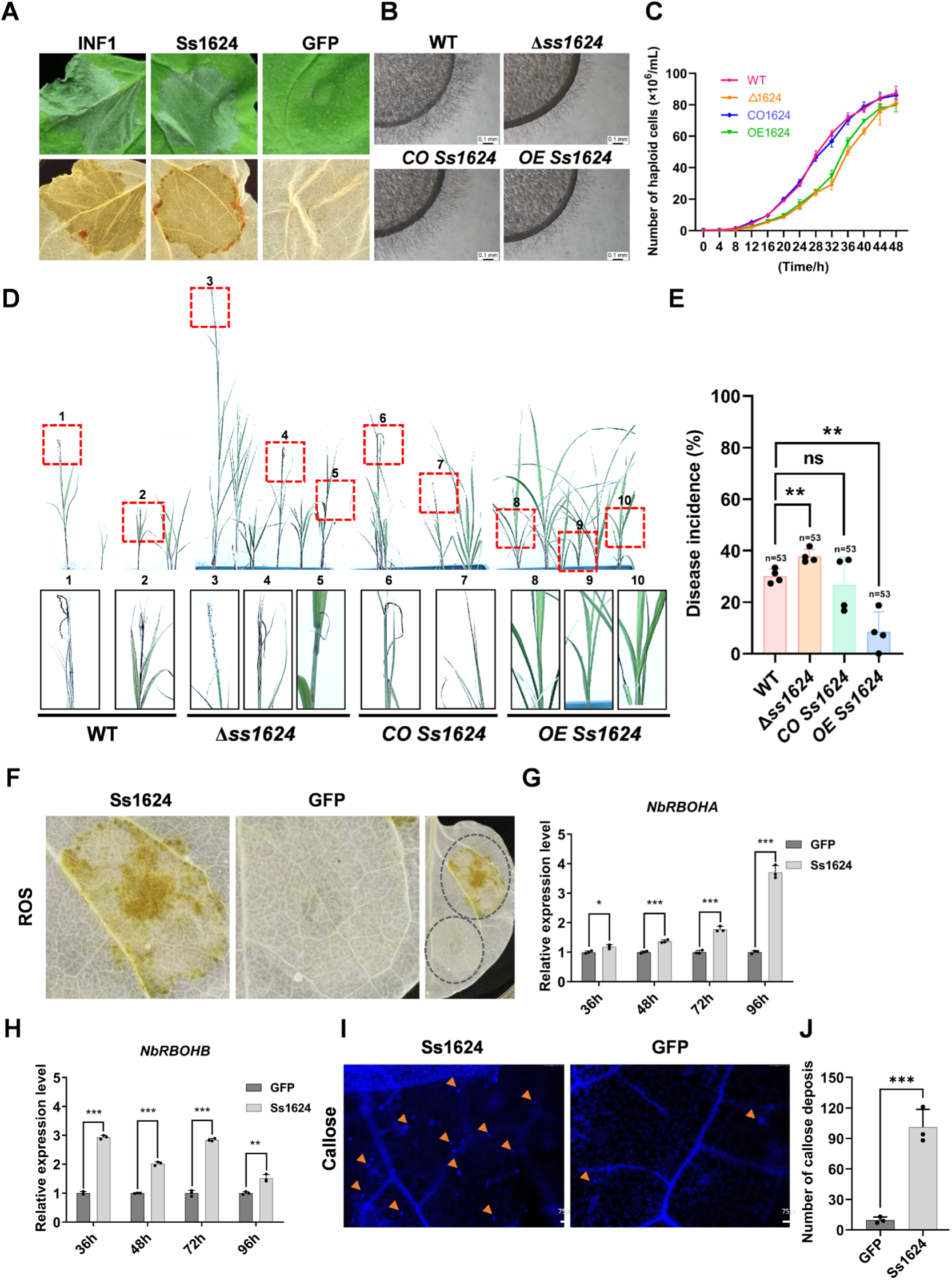
The role of Ss1624 in regulating plant cell death, fungal growth, and immune responses. (A) Cell death induction assays were performed using Ss1624 in 5-week-old *N. benthamiana* plants, with INF1 and GFP serving as positive and negative controls, respectively. (B) Hyphal growth of *S. scitamineum* strains. WT, deletion mutant (Δss*1624*), overexpression strain (*OE Ss1624*), and complemented strain (*CO Ss1624*) were cultured on minimal medium (MM) for 24 h. Hyphal development at colony edges was observed under a stereomicroscope. Scale bars, 0.1 mm. (C) Growth dynamics of haploid cells in liquid culture. Growth curves of the indicated strains (WT, purple; Δss*1624*, orange; *OE Ss1624*, green; *CO Ss1624*, blue) were monitored over 48 h, with cell density (×10^6^ cells/mL) recorded at 4 h intervals. Data are presented as mean ± SD from three independent biological replicates. (D) Pathogenicity assays on sugarcane (cultivar ROC22). Seedlings at the 3-4 leaf stage were inoculated via stem injection with compatible haploid mixtures (OD_600_ = 1.0) of the WT, Δss*1624*, *OE Ss1624*, or *CO Ss1624* strains and maintained under greenhouse conditions for 120-180 days. (E) Disease incidence was calculated as the percentage of symptomatic plants relative to the total number of inoculated plants. (F) Reactive oxygen species (ROS) accumulation following expression of Ss1624. Leaves expressing Ss1624 or GFP (control) were collected at 2 dpi and analyzed for ROS-associated staining. (**G** and **H**) Induction of respiratory burst oxidase homolog genes. Relative transcript levels of *NbRbohA* and *NbRbohB* were quantified by qRT-PCR at 36, 48, 72, and 96 hpi following expression of Ss1624 or GFP. Expression was normalized to *NbEF1α*. (I) Callose deposition induced by Ss1624. Leaves expressing Ss1624 or GFP were harvested at 2 dpi and stained with aniline blue. Callose deposits were visualized under a fluorescence microscope. Scale bars: 75 μm. (J) Quantification of callose deposition. Data are presented as mean ± SD (n = 3). Asterisks indicate significant differences relative to the GFP control (\*\*\**, P* < 0.001; Student’s *t* test).

To assess its biological role, we generated *S. scitamineum* strains with targeted deletion (Δ*ss1624*), complementation (*CO Ss1624*), and overexpression (*OE Ss1624*) of the effector (**Supplemental Figures 2A and 2B**). Under standard *in vitro* growth conditions, these strains exhibited minor differences in colony morphology and growth rates on artificial medium (**Figures 1B and 1C**). Both Δss1624 and *OE Ss1624* strains produced less dense aerial hyphae in mating assays than WT (**Supplemental Figure 2C**), suggesting that appropriate Ss1624 dosage contributes to filamentous growth or mating. By contrast, the mutants did not show consistent differences from WT across the tested osmotic, oxidative, metal-ion and cell-wall stresses (**Supplemental Figures 2D and 2E**).

In sugarcane pathogenicity assays, Δss1624 caused more severe disease than WT, whereas OE Ss1624 was attenuated (**Figures 1D and 1E**). These reciprocal phenotypes indicate that Ss1624 abundance is inversely associated with virulence under the tested conditions. Because Ss1624 overexpression attenuated smut virulence and Ss1624 triggered cell death in *N. benthamiana*, we tested whether it elicited additional immune-associated outputs. Ss1624 expression increased ROS accumulation and callose deposition (**Figures 1F-1J**) and induced defense-marker transcripts associated with salicylic acid, jasmonic acid, ethylene and PTI-related responses (**Supplemental Figures 3A and 3B**). These responses show that Ss1624 is immunogenic in *N. benthamiana*. We therefore used Ss1624-triggered cell death and associated defense outputs as readouts to identify host components required for this response, without assigning the response exclusively to the classical PTI or ETI category.

### Identification of RLK15 as an Ss1624-interacting receptor kinase

To identify host proteins associated with Ss1624, we performed a yeast two-hybrid (Y2H) screen using mature Ss1624 as bait against a cDNA library prepared from *S. scitamineum*-infected sugarcane. RLK15 was recovered as an interacting protein. RLK15 is a CrRLK1L-subfamily receptor-like kinase which contains an extracellular malectin-like domain (MLD), a single-pass transmembrane region, and an intracellular serine/threonine kinase domain. Domain-mapping Y2H assays detected Ss1624 association with both the isolated MLD and the isolated intracellular kinase domain (**Figure 2A**).

**Figure 2.**
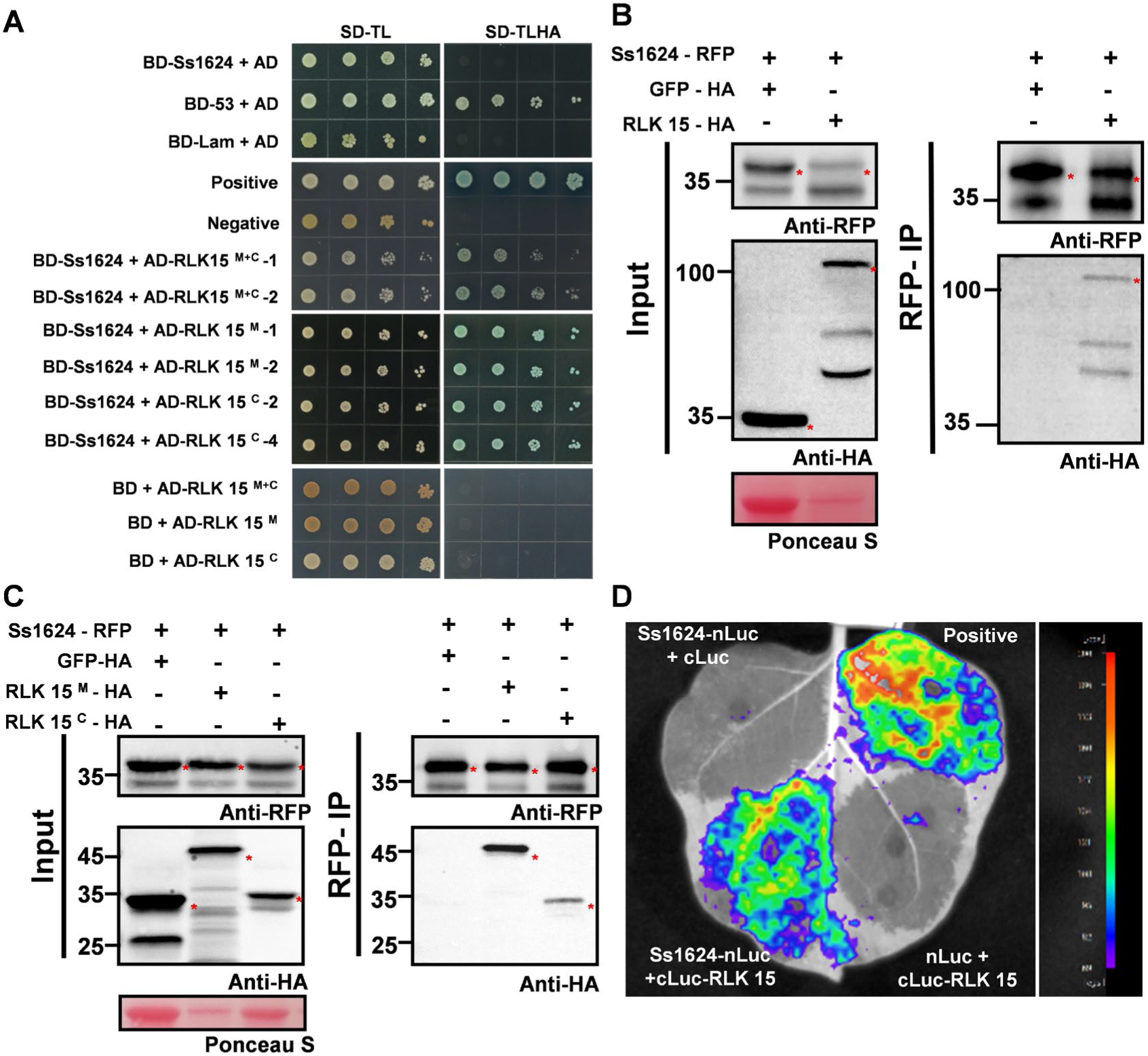
Ss1624 interacts with the receptor-like kinase RLK15. **(A)** Yeast two-hybrid assays demonstrating interaction between Ss1624 and RLK15. Ss1624 was tested for interaction with the RLK15 ^M+C^ domain (lacking the transmembrane region), as well as the individual extracellular (RLK15 ^M^) and intracellular (RLK15 ^C^) domains. Yeast transformants were plated on non-selective medium (SD/- T/-L) to confirm viability and on selective medium (SD/-T/-L/-H/-A) to assess protein-protein interaction. Positive (BD-53 + AD-T) and negative (BD-Lam + AD-T) controls are shown. (**B** and **C**) Co-immunoprecipitation (Co-IP) assays validating the interaction. Protein extracts from leaves co-expressing Ss1624-RFP with GFP-HA, RLK15-HA, RLK15 ^M^-HA, or RLK15 ^C^-HA at 2 dpi were subjected to immunoprecipitation using anti-RFP beads, followed by Western blotting with anti-HA antibody. Ponceau S staining of PVDF membranes was used as a loading control. (**D**) Split-luciferase complementation (SLC) assay confirming interaction in planta. Strong luminescence signals were observed in *N. benthamiana* leaves co-expressing Ss1624-nLuc- and cLuc-RLK15.

We validated these physical interactions *in planta* via co-immunoprecipitation (Co-IP) and split-luciferase complementation (SLC) assays in *N. benthamiana*. Total protein extracts from the leaves co-expressing Ss1624-RFP and full-length RLK15-HA were immunoprecipitated using RFP-Trap magnetic agarose beads. Subsequent anti-HA immunoblotting successfully detected RLK15-HA in the precipitated fraction, confirming the interaction *in planta* (**Fig. 2B**). No interaction was detected in negative controls co-expressing Ss1624-RFP with GFP-HA. To verify the dual-domain binding, further Co-IP assays demonstrated Ss1624-RFP co-precipitated with both the isolated extracellular domain (RLK15 ^M^-HA) and the isolated intracellular kinase domain (RLK15 ^C^-HA) (**Fig. 2C**). In complementary SLC assays, co-expressing Ss1624-nLuc with cLuc-RLK15 reconstituted a strong luminescent signal (**Fig. 2D**). Together, these independent assays establish a physical association between Ss1624 and both domains of RLK15 in planta.

### RLK15 mediates Ss1624-triggered immune cell death in planta

To explore whether related RLK15 homologs in the dicot *N. benthamiana* respond to Ss1624, we identified the two most similar *N. benthamiana* genes, *NbRLK15-1* and *NbRLK15-2* (**Supplemental Figure 4, Supplemental Table 1**). Y2H assays confirmed that Ss1624 also interacts with the intracellular kinase domains of NbRLK15-1 and NbRLK15-2 (**Supplemental Figure 5**). We then tested whether RLK15 itself is required for Ss1624-triggered responses. Simultaneous silencing of *NbRLK15-1* and *NbRLK15-2* significantly suppressed Ss1624-triggered cell death (**Figures 3A-3C, Supplemental Figure 6**). These results together indicate that Ss1624 triggers cell death through an RLK15-dependent signaling pathway.

**Figure 3.**
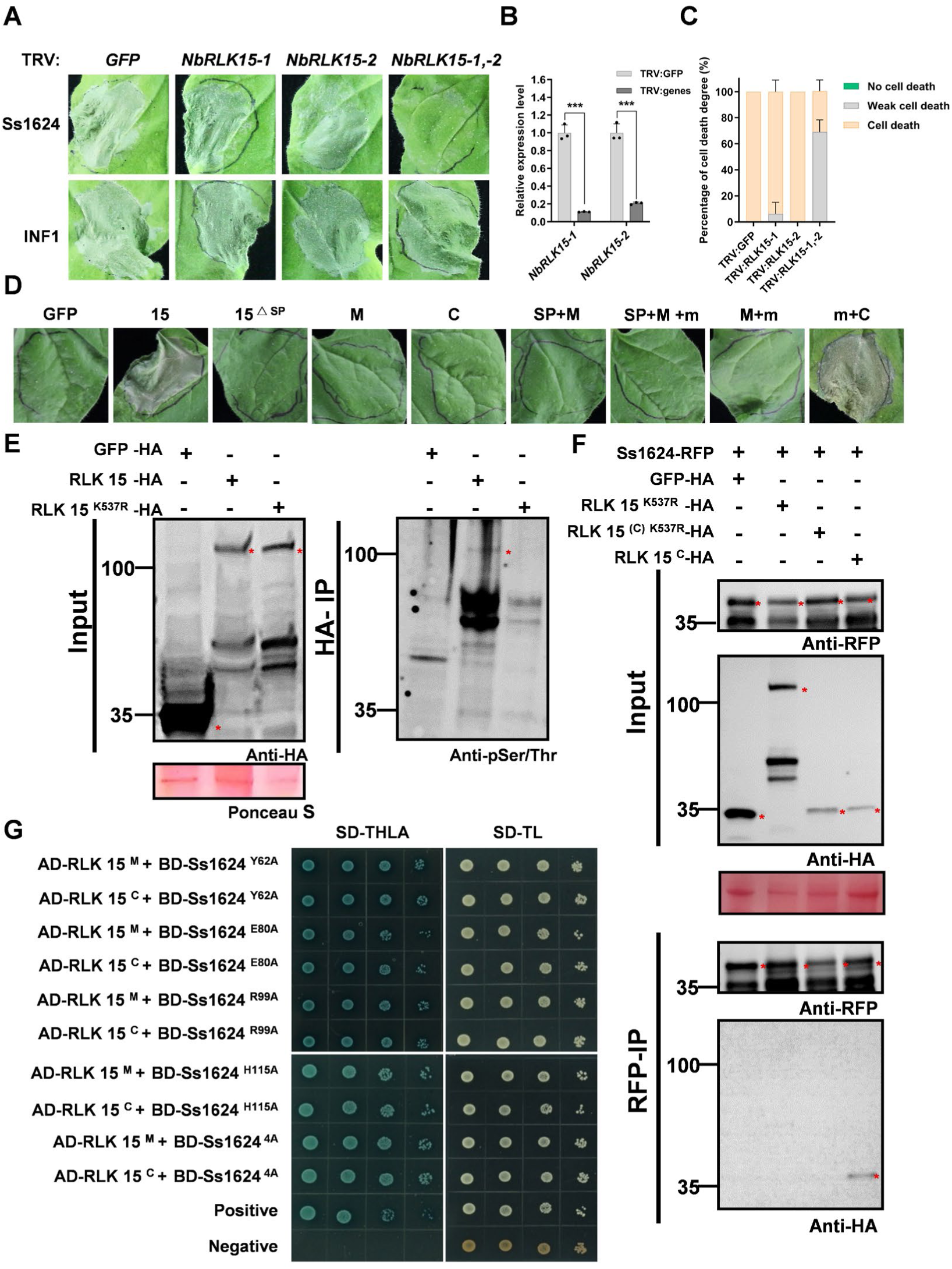
Ss1624-induced cell death mediated by RLK15 and the interaction between RLK15 and Ss1624 depend on the specific domains and kinase activity of RLK15, but not on Ss1624 RNase activity. (A) Leaf phenotypes of *N. benthamiana* at 3 dpi following Ss1624 or GFP expression. VIGS was used to target *NbRLK15* homologs (TRV:*NbRLK15-1*, TRV:*NbRLK15-2*, or both), with TRV:*GFP* serving as a control. (B) qRT-PCR was performed to assess the efficiency of simultaneous silencing of *NbRLK15-1* and *NbRLK15-2* in VIGS-treated plants. Data are shown as mean ± SD (n = 3), with significant differences indicated (***, *P* < 0.001; Student’s *t* test). (C) Quantification of cell death phenotypes induced by Ss1624, as shown in (d). Data are presented as the percentage of leaves exhibiting no, weak, or strong cell death (mean ± SD, n = 8). (D) Membrane localization signal and intracellular kinase domain are required for RLK15 to induce cell death.

### Membrane targeting and kinase activity determine RLK15 immune activity

To define the minimal RLK15 structural requirements necessary for immune activation, we performed a systematic structure-function analysis by transiently expressing deletion and synthetic fusion constructs in *N. benthamiana* (**Figure 3D, Supplemental Figure 7**). While full-length RLK15 consistently induced strong macroscopic cell death, isolated extracellular MLD (M) or intracellular kinase domain (C) alone failed to elicit visible responses. Furthermore, deleting the native signal peptide completely abolished cell death-inducing activity, indicating proper subcellular routing is essential.

Fusing the RLK15 membrane-targeting sequence (m) to the isolated intracellular kinase domain (m + C) rescued cell-death activity. By contrast, constructs that retained or retargeted the extracellular region without the kinase domain (SP + M, M + m and SP + M + m) did not trigger visible cell death (**Supplemental Figure 8)**. Thus, a membrane-tethered RLK15 kinase domain is sufficient to trigger this overexpression-associated response, whereas the kinase domain without membrane targeting is not.

### Kinase activity of RLK15 is essential for Ss1624 binding

We next asked whether the catalytic competence of RLK15 influences its association with Ss1624. Immunoprecipitated 3×HA-RLK15 carried a phospho-Ser/Thr signal that was absent from the GFP control (**Figure 3E**). Sequence alignment with the *Arabidopsis* CrRLK1L receptor FERONIA identified K537 as a conserved lysine required for kinase activity (Haruta *et al*., 2018) (**Supplemental Figure 9, Supplemental Table 2**). Mutating this residue to arginine (K537R) completely abolished *in vivo* autophosphorylation, generating a verified kinase-dead mutant (**Figure 3E**).

To determine whether catalytic activity is necessary for effector perception, we repeated Co-IP assays using this kinase-dead variant. Remarkably, the RLK15 ^K537R^ mutant failed to co-precipitate with Ss1624-RFP, demonstrating that an active, phosphorylation-competent kinase domain is structurally required to form a stable effector-receptor complex *in planta* (**Figure 3F**). Y2H assays independently corroborated this functional requirement, where the K537R mutation similarly disrupted binding (**Supplemental Figure 10**). These results indicate that kinase activity is a structural prerequisite for Ss1624 binding, rather than a downstream consequence of it. Crucially, transient expression of either the full-length kinase-dead mutant (RLK15 ^K537R^) or the membrane-targeted kinase-dead mutant (m + C ^K537R^) failed to trigger cell death (**Figure 3F**). Together with the membrane-targeting experiments, these data establish kinase competence and membrane localization as key determinants of RLK15 activity in this assay.

Ss1624 was predicted to be a fungal-type RNase with four highly conserved catalytic-domain residues, Y62, E80, R99 and H115. In vitro, wild-type Ss1624 efficiently degraded yeast total RNA, whereas the Y62A, R99A and H115A mutants lost detectable activity, comparable to the GFP control (**Supplemental Figure 11**), identifying these three residues as catalytically essential. Because Ss1624 interacts with RLK15, we tested whether this interaction depends on effector’s catalytic activity: in yeast two-hybrid assays, both RLK15 domains (AD-RLK15 M and AD-RLK15 C) bound wild-type Ss1624 (BD-Ss1624), all four single mutants and the quadruple mutant (4A), with blue colonies on SD-TLHA/X-α-gal in every combination (**Figure 3G**). Catalytic activity is therefore dispensable for the Ss1624–RLK15 interaction.

### Ss1624 induces the expression of RLK15 and enhances its autophosphorylation

Given the 60–120-day field disease cycle of sugarcane smut, we monitored transcript levels in *S. scitamineum*-inoculated sugarcane from 2 to 28 days post-inoculation (dpi) to reveal the early temporal dynamics of effector and receptor expression. Time-course qRT-PCR revealed that *Ss1624* transcripts initially declined during early biotrophic establishment (2-4 dpi), followed by a pronounced, sustained increase during later colonization stages. Host *RLK15* expression profiles closely mirrored this pattern, exhibiting simultaneous suppression followed by strong induction (**Figure 4A**).

**Figure 4.**
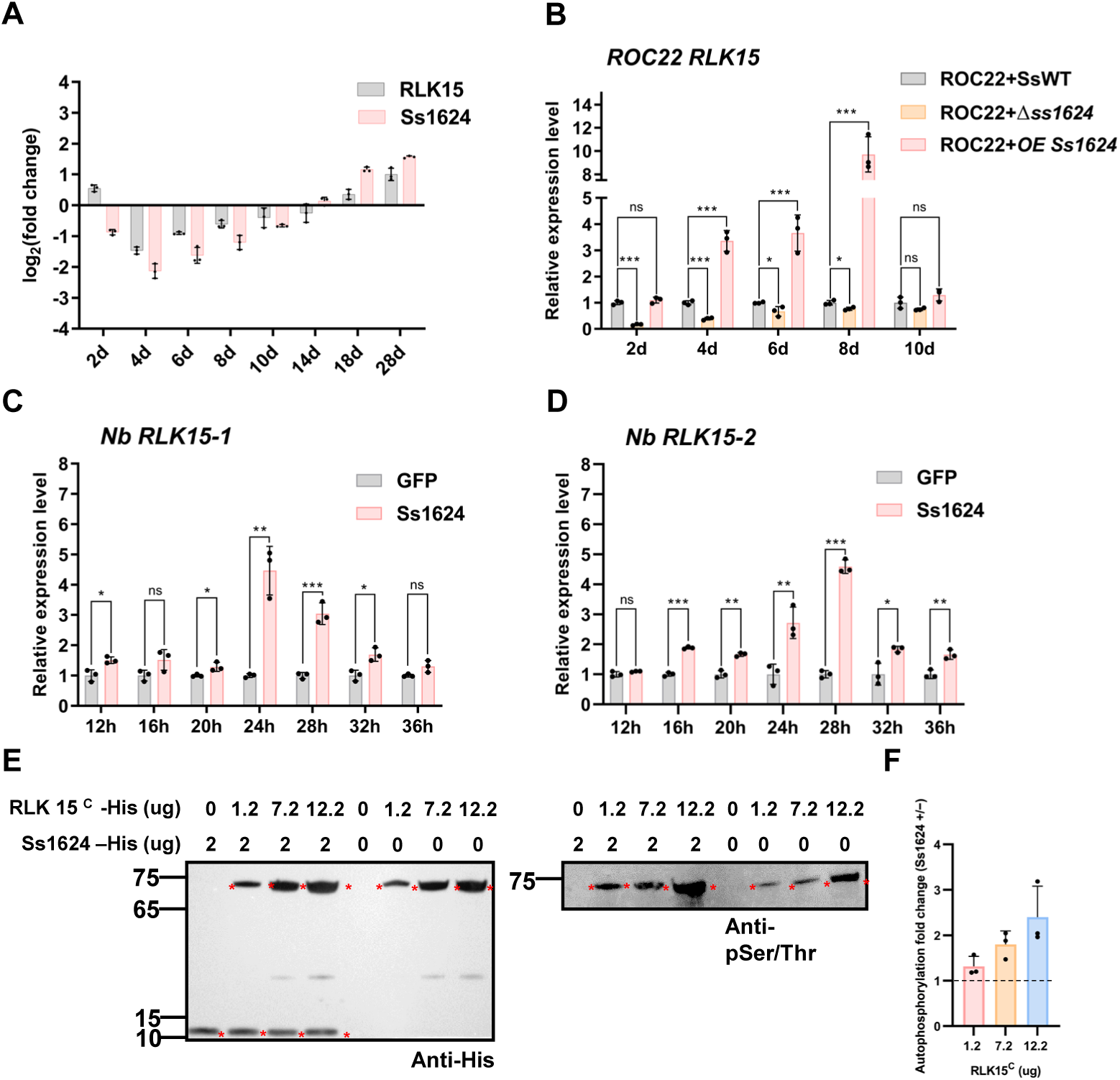
Ss1624 induces RLK15 expression and enhances its autophosphorylation. (A) Temporal expression profile of *Ss1624* and *RLK15* in sugarcane cultivar ROC22 during *S. scitamineum* infection. Transcript abundance was quantified at indicated time points post-inoculation and data are presented as log_2_(fold change) relative to the mock control. (B) Dependence of RLK15 induction on Ss1624. *RLK15* transcript levels were measured in plants inoculated with WT, Δ*ss1624*, or *OE Ss1624* strains, revealing enhanced and sustained expression in the *OE Ss1624* treatment. (**C** and **D**) Induction of the *RLK15* homolog in *N. benthamiana*. Relative expression levels of *NbRLK15-1* and *NbRLK15-2* were quantified following transient expression of Ss1624. Data represent mean ± SD (n = 3), with statistical significance indicated (*, *P* < 0.05; **, *P* < 0.01; ***, *P* < 0.001; ns, not significant; Student’s *t* test). (E) Ss1624 enhances autophosphorylation of the RLK15 ^C^-terminal domain *in vitro*. Kinase reactions were performed using increasing amounts of RLK15 ^C^ (0, 1.2, 7.2, and 12.2 μg) in the presence or absence of Ss1624 (2 μg). Phosphorylation levels were assessed by Western blot using an anti-phosphoserine/threonine antibody, with anti-His serving as a loading control. (F) Quantitative statistical comparison of RLK15 ^C^ autophosphorylation in (E). Band intensities were quantified with ImageJ. The total-protein-corrected phosphorylation signal was obtained by dividing the net intensity of the phosphorylated RLK1 ^C^ band by the net intensity of the anti-His (total protein) band. The relative phosphorylation level was expressed relative to the mean corrected ratio of the Ss1624 (0 μg) group at 1.2 μg RLK15 ^C^, which was set to 1. The fold change was then calculated as the relative phosphorylation level with Ss1624 divided by that without Ss1624 at the same RLK15 ^C^ amount. Data are mean ± SD of three independent biological replicate. Constructs corresponding to GFP (negative control), full-length RLK15 (15), RLK15 lacking the SP (15 ^ΔSP^), isolated extracellular domain (M), isolated intracellular kinase domain (C), extracellular domain fused to a SP and membrane-targeting sequence (SP + M + m), extracellular domain with membrane localization signal (M + m), and intracellular domain with membrane localization signal (m + C) were transiently expressed in *N. benthamiana* leaves via *Agrobacterium* infiltration. Cell death was observed only in constructs containing a functional intracellular kinase domain together with membrane targeting (m + C), indicating that both kinase activity and membrane localization are required. (E) In vivo autophosphorylation activity of RLK15. The WT protein exhibits autophosphorylation, whereas the kinase-deficient mutant (K537R) abolishes this activity. (F) Kinase activity of RLK15 is required for interaction with Ss1624. Co-immunoprecipitation assay demonstrates that the K537R kinase-dead mutation disrupts the binding between RLK15 and Ss1624. (G) The RNase catalytic activity of Ss1624 does not affect its interaction with RLK15. Yeast cells co-expressing AD-RLK15 ^M^ or AD-RLK15 ^C^ with the Ss1624 catalytic-site single mutants (BD-1624 ^Y62A^, BD-1624 ^E80A^, BD- 1624 ^R99A^, BD-1624 ^H115A^) or the quadruple mutant (BD-1624 ^4A^) grew on selective medium (SD/-T/-L/-H/-A).

Given this coordinated expression pattern, we next asked whether *Ss1624* expression is associated with *RLK15* accumulation. Profiling *RLK15* in sugarcane inoculated with *S. scitamineum* mutant strains showed that, relative to WT infection, *RLK15* accumulation was significantly reduced at all time points in Δ*ss1624*-infected plants. Conversely, *OE Ss1624* infection provoked markedly elevated *RLK15* expression, peaking strongly at 8 dpi (**Figure 4B**). This tightly correlated expression suggests a positive regulatory feedback loop wherein Ss1624 accumulation drives transcriptional induction of its associated receptor.

To test this hypothesis directly, we examined whether this regulation is conserved in a heterologous system: in *N. benthamiana,* transient Ss1624 expression strongly induced transcription of both *NbRLK15* homologs (**Figures 4C and 4D**). Furthermore, *in vitro* phosphorylation assays using recombinant RLK15 intracellular domain (RLK15 ^C^) purified from *Escherichia coli* demonstrated that incubation with purified Ss1624 significantly enhanced receptor autophosphorylation relative to controls (**Figure 4E and 4F, Supplemental Figure 12**). These findings suggest a dynamic recognition model in which Ss1624 treatment or expression both elevates RLK15 abundance via transcriptional induction and directly activates the existing RLK15 protein by enhancing its autophosphorylation.

### RLK15 plays a conserved role in immunity and confers resistance to sheath blight, rice blast and bacterial blight

Finally, having established RLK15 as a specific immune receptor, we evaluated its broader translational potential by characterizing its ortholog (OsRLK15, LOC_Os07g05370) in the major global staple crop rice (*Oryza sativa* L. *japonica* cv Zhonghua 11, ZH11), because the difficulties of genetic transformation in sugarcane and the limitations of the specific disease types affecting sugarcane precluded validation of RLK15 function in its native host. We generated CRISPR/Cas9 osrlk15 knockout plants (**Supplemental Figure 13**) and stable OsRLK15 overexpression lines driven by the *CsVMV* promoter from cassava vein mosaic virus. These genotypes were challenged with the sheath blight pathogen *Rhizoctonia solani*, the fungal blast pathogen *Magnaporthe oryzae* and the bacterial blight pathogen *Xanthomonas oryzae* pv. *Oryzae* (*Xoo*).

Following *R. solani* and *M. oryzae* inoculation, *osrlk15* knockout plants developed significantly more extensive and severe disease lesions than WT plants, indicating enhanced susceptibility. Conversely, *OsRLK15* overexpression lines exhibited increased resistance, characterized by more restricted, smaller lesions (**Figures 5A-5D**). A similar direction of effect was observed after *Xoo* inoculation: *osrlk15* leaves developed longer lesions and overexpression lines shorter lesions than WT (**Figures 5E and 5F**). Under the field conditions used for agronomic phenotyping, the measured traits of *OsRLK15* overexpression lines, including grain dimensions, thousand-grain weight, panicle length, kernel number, seed-setting rate and tiller number, did not differ detectably from WT (**Supplemental Figures 14A-14I**). Taken together, these results demonstrate that OsRLK15 positively regulates rice resistance to tested pathogens, with no detectable agronomic penalty under the conditions examined.

**Figure 5.**
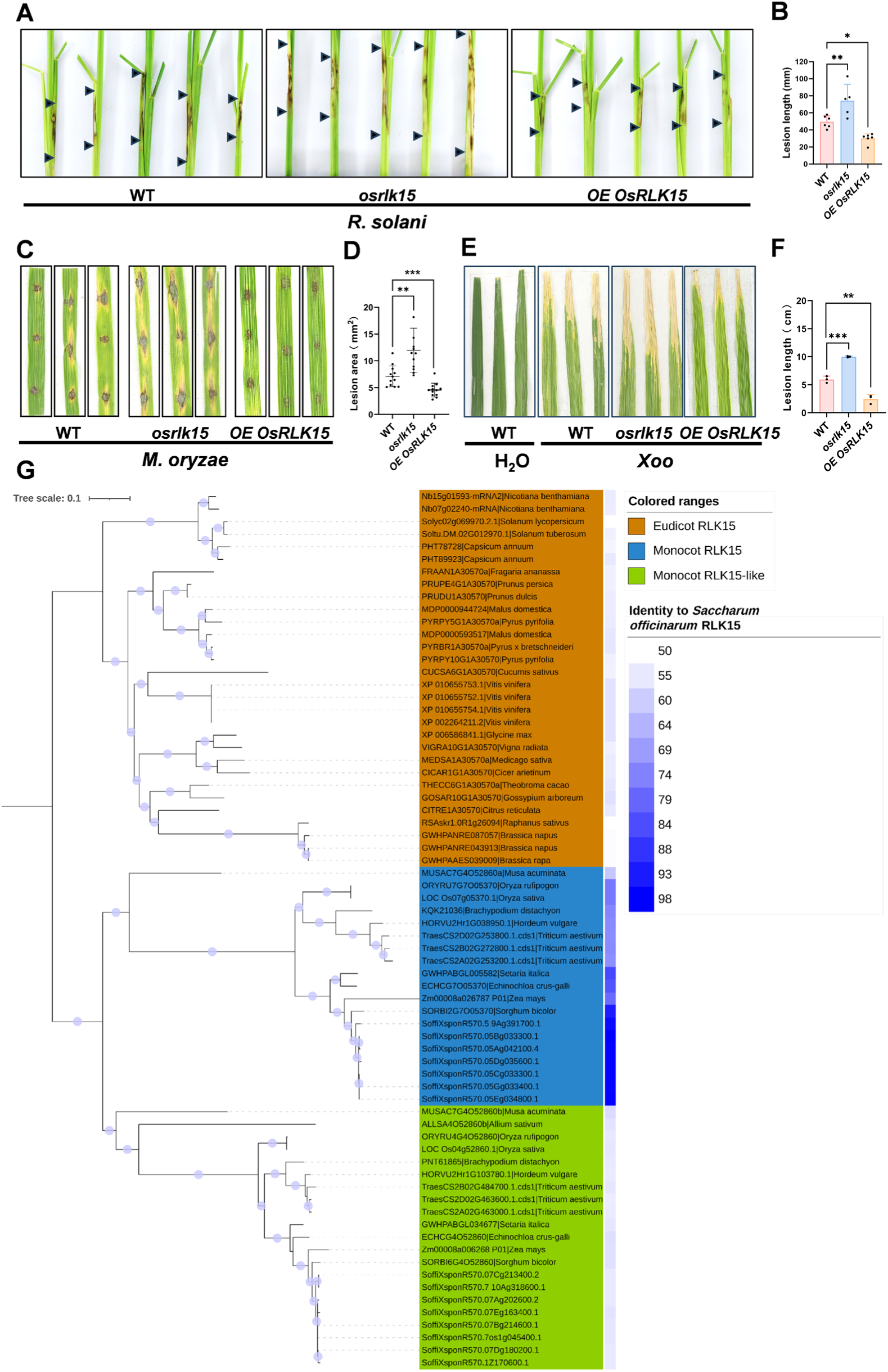
OsRLK15 is required for rice immunity against sheath blight, rice blast and bacterial blight. (A) Representative phenotypes of rice sheath blight from WT (ZH11), *osrlk15*, and *OE OsRLK15* plants at 7 dpi with *R. solani*. Black arrowheads indicate the inoculation sites and typical lesion development. **(B)** Quantitative analysis of lesion lengths. Data are presented as mean ± SD (n ≥ 5). One-way ANOVA followed by Tukey’s honestly significant difference (HSD) test was performed to determine significant differences among multiple groups. Statistical significance was set at *P* < 0.05 (*, *P* < 0.05; **, *P* < 0.01; Student’s *t* test). (C) Leaf phenotypes of rice plants following inoculation with *M. oryzae* at 7 dpi. WT (ZH11), *osrlk15*, and *OE OsRLK15* plants were compared. Lesion areas were markedly larger in *osrlk15* mutants and reduced in *OE OsRLK15* plants relative to wild type. (D) Quantification of lesion areas shown in (c). Data are presented as mean ± SD (n = 12). Significant differences relative to the WT control are indicated (**, *P* < 0.01; ***, *P* < 0.001; one-way ANOVA followed by Tukey’s HSD test). (E) Disease symptoms in rice plants inoculated with *Xoo* at 14 dpi. Leaves from WT ZH11, *osrlk15*, and *OE OsRLK15* plants were examined, with water-inoculated leaves serving as a negative control. The *osrlk15* mutant showed increased susceptibility, whereas *OE OsRLK15* plants displayed enhanced resistance. (F) Quantification of lesion lengths caused by *Xoo*. Data are presented as mean ± SD (n ≥ 3). Significant differences relative to WT are indicated (**, *P* < 0.01; ***, *P* < 0.001; one-way ANOVA followed by Tukey’s HSD test). (G) Phylogenetic analysis of the plant RLK15 (CrRLK1L) subfamily. Phylogenetic reconstruction of the CrRLK1L family across angiosperms was conducted using the metaRLK database. The analysis revealed that RLK15 belongs to a well-supported monophyletic clade originating from a single ancestral gene. A distinct monocot-wide duplication event was identified, resulting in single-copy orthologs in diploid crops and multiple copies in polyploids. Remarkably, RLK15 exhibits high sequence conservation with its homologs, displaying amino acid sequence identities ranging from 50% to 98%.

We next examined the evolutionary distribution of RLK15-related receptors. Phylogenetic analysis of the CrRLK1L receptor-like kinase family across monocots and eudicots using the metaRLK database (Liu *et al*., 2024) (**supplementary data 1**), which revealed that RLK15 belongs to a well-supported clade descending from a single ancestral gene present in the common ancestor of all angiosperms (**Figure 5G**). Within this clade, we identified a distinct monocot-wide duplication event, which yielded single-copy orthologs in most diploid crops, such as rice and maize, and multiple copies in complex polyploids, including wheat and sugarcane. Strikingly, RLK15 shares sequence conservation with its homologs across these diverse plant species, with amino acid sequence identity ranging from 50% to 98%. To clarify the precise evolutionary placement of RLK15 within the CrRLK1L superfamily, we built a maximum-likelihood tree using ROC22-RLK15 and key *Arabidopsis thaliana* CrRLK1L members (**Supplemental Figure 15**). The results show that RLK15 forms a tight cluster with AtHERK2 (100% bootstrap support), consistent with the striking sequence conservation (50–98% identity) observed between RLK15 and its homologs across diverse angiosperm lineages.

In parallel, we evaluated the evolutionary conservation of the Ss1624 effector across the fungal kingdom. Interrogation of the OrthoDB v12.2 database showed that Ss1624 belongs to the widely distributed “400260at4751” fungi-level orthogroup (**Supplemental Figure 16**). Subsequent phylogenetic reconstruction of this orthogroup demonstrated that numerous diverse fungal pathogens carry functional homologs of Ss1624. Notably, the effector exhibits exceptional sequence conservation among species within the *Ustilago* and *Sporisorium* genera.

The co-conservation of both the effector and its cognate receptor across divergent plant and fungal lineages suggests that the Ss1624–RLK15 recognition module may represent a broadly conserved effector–receptor interaction in plant immunity. One that could operate in crop–pathogen systems beyond those tested here.

## Discussion

Pathogens secrete effectors to suppress plant immunity, but these can be perceived by host receptors, creating reciprocal evolutionary pressure that drives effector diversification. Our results identify Ss1624 as an RNase-type fungal effector that is simultaneously enzymatically active and immunogenic, and connect its recognition-associated phenotype to the CrRLK1L receptor kinase RLK15. Ss1624 induces ROS accumulation, callose deposition, defense-associated transcription and cell death in *N. benthamiana*, while RLK15 homologues are genetically required for the cell-death response. Together with the biochemical observation that Ss1624 enhances RLK15 kinase-domain autophosphorylation in vitro, these findings support a model in which a pathogen-derived RNase is coupled to activation of a malectin-like receptor kinase (**Figure 6**).

**Figure 6.**
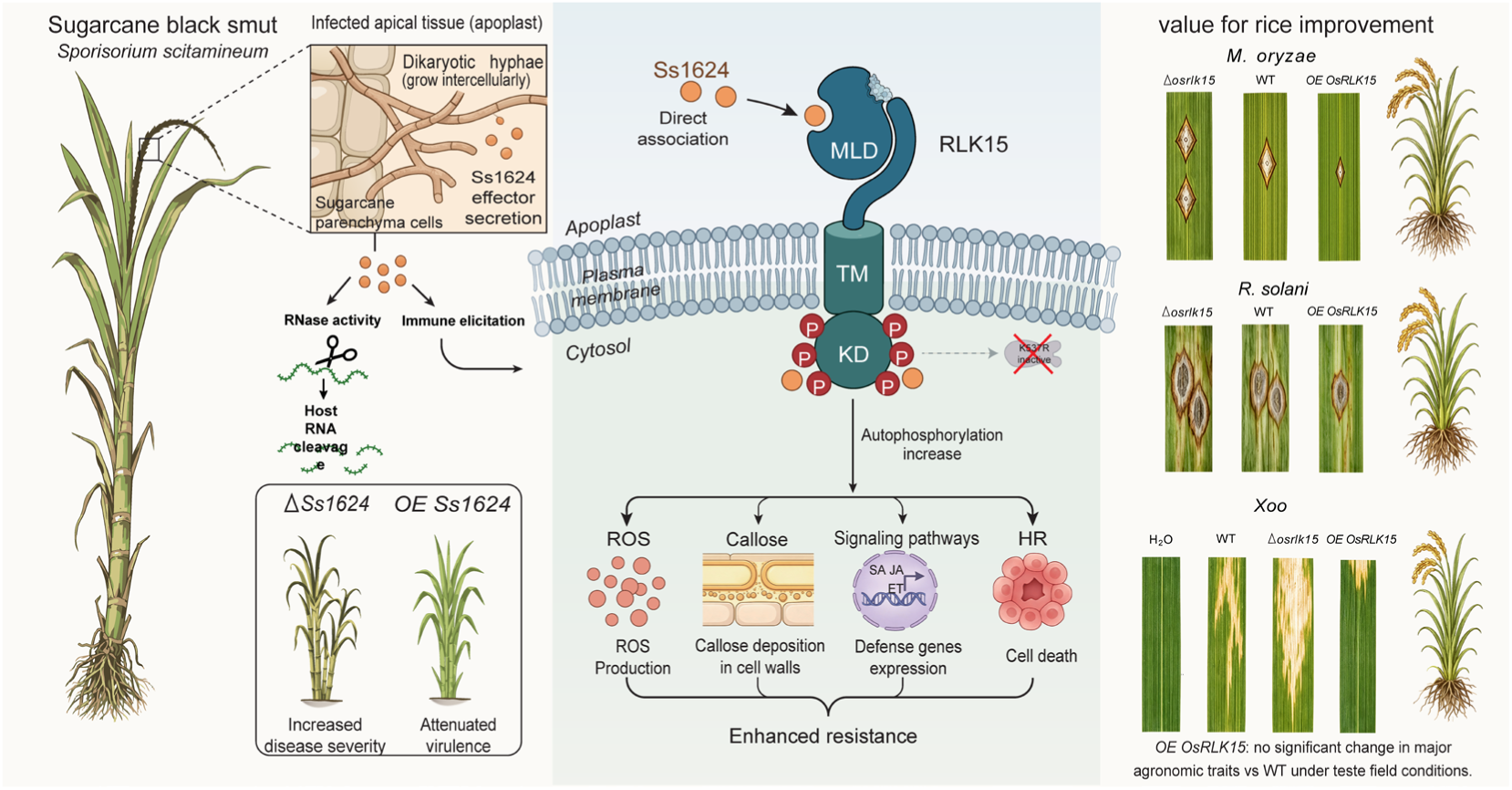
Schematic model for the role of Ss1624 and RLK15 in plant immunity. The fungal effector Ss1624, a secreted ribonuclease can localize to the plant apoplast, degrades host RNA while also functioning as an immune elicitor. At the plasma membrane, the CrRLK1L family receptor-like kinase RLK15 recognizes and interacts with Ss1624. This interaction promotes RLK15 autophosphorylation and activates downstream signaling cascades. As a consequence, immune responses are induced, including reactive oxygen species production, callose deposition, and transcriptional activation of defense pathways mediated by salicylic acid (SA), jasmonic acid (JA), and ethylene (ET), ultimately contributing to enhanced disease resistance.

Ss1624 exemplifies the delicate balance that biotrophic fungi must maintain during host colonization. Deletion of Ss1624 resulted in heightened fungal virulence and exacerbated disease symptoms, whereas its constitutive overexpression severely attenuated disease progression (**Figures 1D and 1E**). This contrasts with the previously characterized RNase effectors such as Fg12 (*F. graminearum*), FoRnt2 (*F. oxysporum*), SRE1 (*S. turcica*), and Nuc1/Nuc2 (*U. maydis*), whose loss typically lead to reduced virulence (Mukherjee *et al*., 2020; Yang *et al*., 2021; He *et al*., 2022; Qian *et al*., 2022). Instead, Ss1624 resembles the other group of RNase effectors such as VdRTX1 (*V. dahliae*) and SRN1/SRN2 (*C. orbiculare*), where loss of function increases the fungal virulence (Yin *et al*., 2022; Kumakura *et al*., 2024). In contrast, overexpression of the RNase effector Ribo1 (*U. maydis*) reduces virulence, whereas the SG200 Δ*ribo1* mutant shows no significant difference in virulence compared with the wild type. (Ökmen *et al*., 2023) These observations highlight the functional diversity of RNase effectors and their potential to act as either virulence factors or immune elicitors. These RNase effectors might facilitate initial virulence through their RNA-degrading capability, which either disrupts host cellular programming or silences defense-related transcripts or both during the early stages of infection. However, its over-accumulation may incur an evolutionary disadvantage by rendering the pathogen detectable to the host immune system. Consistent with this, *Ss1624* exhibited a highly coordinated, biphasic expression pattern characterized by initial suppression followed by late-stage induction, suggesting that the fungus tightly titrates its secretion to remain below the host’s detection threshold during early biotrophic establishment.

Beyond virulence modulation, Ss1624 also contributes to core fungal physiology, as its deletion impaired mating or hyphal development (**Supplemental Figure 2C**). This distinguishes Ss1624 from many RNase effectors that do not affect fungal growth (Yang *et al*., 2021; Qian *et al*., 2022), and supports an expanded view of effector biology in which effectors can simultaneously participate in pathogen fitness and host manipulation (de Guillen *et al*., 2025; Ma, T *et al*., 2025). The biphasic expression pattern, that is, initial suppression followed by late-stage induction, resembles regulatory mechanisms described in other pathosystems, including chromatin-mediated control of the *F. graminearum* xylanase BCG1 (Zhao *et al*., 2024) and suppression of immunogenic flagellar components in *Pseudomonas syringae* via AlgU (Pel *et al*., 2014; Bao *et al*., 2020). These parallels suggest that temporal regulation of immunogenic effectors may represent a conserved strategy among pathogens to balance virulence and immune evasion.

To identify the host receptor for Ss1624, a yeast-two-hybrid screen identified RLK15, a receptor whose expression is co-regulated with the effector during infection (**Figure 4A**). RLK15 belongs to the *Catharanthus roseus* receptor-like kinase 1-like (CrRLK1L) subfamily of receptor-like kinases. Members of the CrRLK1L family, such as FERONIA (FER), THESEUS1 (THE1), and ANXUR1/2 (ANX1/2) in *Arabidopsis*, are receptor kinases characterized by an extracellular malectin-like domain (MLD).(Cheung & Wu, 2011; Franck *et al*., 2018; Solis-Miranda & Quinto, 2021) They have long been established as sensors of cell wall integrity, regulators of polarized cell growth, and mediators of fertilization, primarily by perceiving endogenous Rapid Alkalinization Factor (RALF) peptides via their MLD.(Escobar-Restrepo *et al*., 2007; Feng *et al*., 2018; Xiao *et al*., 2019; Zwartkruis *et al*., 2026) In plant immunity, FERONIA exhibits highly context-dependent dual functions. It acts as a molecular scaffold that facilitates the ligand-induced assembly of immune receptors FLS2 and EFR with their co-receptor BAK1, thereby positively regulating pattern-triggered immunity (PTI).(Stegmann *et al*., 2017) Conversely, upon binding endogenous RALF23, FER inhibits FLS2-BAK1 complex formation and negatively regulates immune signaling (Stegmann et al., 2017). This inhibition is finely tuned by RALF23 concentration: low concentrations promote receptor complex assembly, whereas high concentrations suppress it.(Chen *et al*., 2024) THESEUS1 (THE1) functions as a RALF34 receptor in *Arabidopsis* and mediates the response of growing plant cells to perturbation of cellulose synthesis, acting as a cell wall integrity sensor.(Gonneau *et al*., 2018) ANXUR1/2 (ANX1/2) not only negatively regulates PTI by competitively binding BAK1 to impede FLS2-BAK1 complex formation, but also suppresses ETI by promoting RPS2 degradation and inhibiting NLR-mediated cell death.(Mang *et al*., 2017) Moreover, studies have established the CrRLK1L family as a key target of diverse trans-kingdom pathogens. For instance, *F. oxysporum* secretes the RALF-mimetic peptide F-RALF,(Masachis *et al*., 2016) and root-knot nematodes produce MiRALF1/3.(Zhang *et al*., 2020) Both function by mimicking endogenous RALF peptides to bind the extracellular domain of FER, thereby hijacking host signaling to suppress immunity and promote infection. During Turnip mosaic virus (TuMV) infection, the viral 6K2 protein directly interacts with NbRALF1 and promotes its degradation via the 26S proteasome pathway, thereby facilitating systemic infection.(Rui *et al*., 2025) Despite these advances, no previous study has demonstrated that a CrRLK1L family receptor recognizes a fungal proteinaceous effector. The present study significantly advances this understanding by demonstrating that Ss1624, an RNase effector from *S. scitamineum*, targets and activates RLK15.

First, our findings extend the functional landscape of the CrRLK1L family in plant immunity. Beyond the well-characterized regulatory roles of FER in modulating PTI signaling through immune receptor complex assembly,(Stegmann *et al*., 2017) we identify RLK15 as a immune receptor capable of recognizing a pathogen-derived effector. This discovery suggests that the involvement of the CrRLK1L family in plant immunity may be more extensive than previously appreciated, with different family members potentially adopting distinct modes of immune function. While FER primarily acts as a signaling scaffold that fine-tunes PTI outputs, RLK15 appears to function as a bona fide recognition receptor that senses pathogen invasion. RLK15 therefore represents a second CrRLK1L family member, beyond FER, with a clearly defined role in plant immunity, broadening our understanding of the functional repertoire of CrRLK1L receptors.

Second, RLK15 recognizes a fungal RNase effector through a mechanism fundamentally distinct from the RALF mimicry strategy employed by previously characterized pathogen effectors. Known pathogen-derived molecules targeting CrRLK1L receptors, such as F-RALF from *F. oxysporum* and MiRALF1/3 from root-knot nematodes, exploit structural mimicry of endogenous RALF peptides to competitively occupy the same ligand-binding site within the MLD.(Masachis *et al*., 2016; Zhang *et al*., 2020) In striking contrast, Ss1624 is a structurally unrelated RNase effector that binds and activates RLK15 via a non-canonical mechanism (**Figure 4E**). To our knowledge, this is the first demonstration that a CrRLK1L family receptor can recognize a fungal RNase effector, establishing a new paradigm for how the CrRLK1L family participates in plant immune recognition.

Third, Ss1624 engages RLK15 through an unprecedented dual-domain interaction, binding both the extracellular MLD and the intracellular kinase domain (**Figures 2A and 2C**). This pattern is exceptionally rare in plant–microbe interaction studies. Notably, this interaction is dependent on receptor kinase activity itself, as the kinase-dead mutation K537R abolished both autophosphorylation and effector binding (**Figures 3B and 3C**), revealing a reciprocal interdependence between receptor activation status and effector engagement. This dual-domain binding pattern suggests that Ss1624 may function as an allosteric modulator, potentially stabilizing an active conformation of RLK15 or promoting its dimerization and trans-phosphorylation to achieve integrated activation of the full-length receptor. This mechanism contrasts sharply with the conventional ligand-receptor paradigm in the CrRLK1L family, where RALF peptides exclusively engage the extracellular MLD to elicit downstream signaling. The biological significance of this interaction is further underscored by the unique virulence phenotype of Ss1624. Unlike previously reported RNase effectors,(Mukherjee *et al*., 2020; Yang *et al*., 2021; He *et al*., 2022; Qian *et al*., 2022) deletion of Ss1624 enhanced fungal virulence, whereas its overexpression attenuated pathogenicity (**Figures 1D and 1E**). Combined with its expression pattern, which was suppressed during early infection and strongly induced at later stages (**Figure 4A**), these results indicate that *S. scitamineum* must finely tune Ss1624 secretion levels. This balancing act between immune evasion and virulence requirements suggests that the RLK15-Ss1624 module represents a carefully balanced molecular arms race. In this arms race, the host has evolved RLK15 as a surveillance receptor, whereas the pathogen deploys a precisely regulated effector to manipulate host immunity.

Phylogenetic analyses trace RLK15 back to a single ancestral gene in early angiosperms, maintaining 50% to 98% sequence identity across modern crops (**Figure 5G**). Concurrently, Ss1624 belongs to a highly conserved fungal orthogroup (OrthoDB: 400260at4751) distributed across numerous pathogenic taxa (**Supplemental Figure 16**). This dual evolutionary conservation, where both the pathogen’s weapon and the host’s surveillance system have remained structurally locked across millions of years, suggests that this interaction represents a putative conserved module in plant-pathogen biology.

Translating our findings from sugarcane to rice, the orthologous receptor OsRLK15 confers strong resistance to the tested pathogens including sheath blight, rice blast and bacterial blight (**Figures 5A-5F**). Notably, key agronomic traits such as tiller number, panicle architecture, and grain yield showed no significant reduction in the overexpression lines (**Supplemental Figure 14**). This indicates that simply increasing RLK15 abundance could prime plants for rapid defense activation without constitutively draining the metabolic resources required for development.

In conclusion, we show that a secreted RNase effector serves as an immune elicitor sensed by the host kinase RLK15 to activate defenses. This reveals a dynamic interplay between pathogen elicitors and host surveillance, framing how RNase effectors are integrated into immune recognition. By linking mechanistic insight with translational potential, the present study advances both fundamental knowledge and practical approaches for improving crop resistance in the context of sustainable agriculture.

## Materials and Methods

### Plant materials and growth conditions

*N. benthamiana* plants were cultivated in plastic pots (7 cm × 7 cm × 8 cm) filled with a 1:3 (v/v) mixture of Miracle-Gro potting soil and vermiculite. Plants were maintained in a controlled glasshouse environment under a 16 h light/8 h dark photoperiod at 25 °C. Two-week-old seedlings were individually transplanted and further grown under identical conditions. Unless otherwise indicated, all experiments were conducted using 4-week-old plants, consistent with previously described protocols (Li, 2011).

Sugarcane (*Saccharum* spp. hybrids) cultivar ROC22 was grown in a greenhouse maintained at 28 °C. Healthy, disease-free plants were selected as seed cane material. Prior to planting, cuttings were surface-treated for 30 min in carbendazim solution prepared by diluting a 50% carbendazim stock 1:1000 (v/v), following established procedures (Guo *et al*., 2024). Plants were subsequently planted in sterilized soil and maintained under standard greenhouse management practices until sampling.

### Strains, plasmids, and culture conditions

*S. scitamineum* strains *MAT-1* and *MAT-2*, isolated from whip-like sori developed on infected sugarcane stem apices collected in Guangdong Province, China, were used as parental strains for genetic manipulation. Fungal cultures were routinely maintained on yeast extract-peptone-sucrose agar (YEPS) plates or in liquid YEPS medium under dark conditions at 28 °C to promote growth and mating.

*E. coli* strain JM109 was cultured in Luria-Bertani (LB) medium at 37 °C and used for plasmid cloning and propagation. *A. tumefaciens* strain GV3101 was maintained in LB medium at 28 °C and employed for plant transient expression assays.

DNA fragments for gene knockout, complementation, and overexpression constructs were amplified using Phanta Max Super-Fidelity DNA Polymerase (Vazyme Biotech, Nanjing, China) from genomic DNA of *S. scitamineum* or *U. maydis*. Coding sequences used for transient expression in *N. benthamiana* were amplified from cDNA prepared from *S. scitamineum*, *U. maydis*, or sugarcane tissues. Primers and plasmid constructs were listed in **Supplemental Tables 3 and 4**.

PCR products were digested with SmaI and cloned into pGR107-HA or pBin-HA/pBin-RFP vectors using the ClonExpress II One Step Cloning Kit (Vazyme Biotech, Nanjing, China). VIGS constructs for *N. benthamiana* were generated using the TRV2 vector backbone. Gene fragments for VIGS were amplified from a *N. benthamiana* cDNA library. All constructs were verified by Sanger sequencing (Tsingke Biotech, Beijing, China) prior to use.

### Construction of deletion mutants, complementation and overexpression strains

Genetic manipulation of *S. scitamineum* mutants was performed using truncated resistance marker fragments coupled with polyethylene glycol (PEG)-mediated protoplast transformation (Chang *et al*., 2019). Deletion of Ss1624 was achieved by homologous recombination using a hygromycin resistance cassette. Approximately 1 kb upstream and downstream flanking sequences of the Ss1624 locus were amplified using primer pairs Ss1624-pro1F/Ss1624::HY-gR and h-YGF::Ss1624/Ss1624-tail, respectively. These fragments were fused with partial hygromycin resistance gene fragments (HY/g and h/YG) via overlap PCR, generating two overlapping constructs (5′Ss1624::HY/g and h/YG::3′Ss1624) containing the hygromycin phosphotransferase (HYG) marker. The resulting recombination cassette replaced the endogenous Ss1624 coding region through homologous recombination following PEG-mediated transformation of fungal protoplasts. Transformants were selected on regeneration medium supplemented with 100 μg/mL hygromycin. Putative deletion mutants were screened by PCR using primers flanking the recombination junctions to confirm correct integration and gene replacement.

For generation of complementation constructs, a zeocin resistance-based strategy was employed. Two truncated, overlapping fragments of the zeocin resistance gene (ZE/o and z/EO) were fused to the upstream and downstream flanking regions of Ss1624 using primer pairs Ss1624-pro1F/Ss1624::ZE-oR and z-EOF::Ss1624/Ss1624-tail, respectively. This generated two overlapping fragments (5′Ss1624::ZE/o and z/EO::3′Ss1624) encompassing the complete zeocin resistance cassette (ZEO). The upstream fusion fragment included the full-length Ss1624 coding sequence driven by its native promoter, allowing restoration of endogenous expression patterns.

For overexpression strains, the native Ss1624 promoter was replaced with the constitutive glyceraldehyde-3-phosphate dehydrogenase (GAP) promoter, while maintaining the same vector backbone and recombination strategy used for complementation constructs. Both complementation and overexpression constructs were co-transformed into the Δss*1624* mutant background via PEG-mediated protoplast transformation. Transformants were selected on regeneration medium containing 100 μg/mL zeocin.

Successful integration of complementation or overexpression constructs was verified by PCR and qPCR using gene-specific primers. Correct expression of Ss*1624* in complemented and overexpression strains was further assessed to confirm restoration or enhancement of transcriptional levels relative to WT strains. All primers used throughout this study are listed in **Supplemental Table 3**.

### Agrobacterium-mediated transient expression of S. scitamineum effectors

Recombinant plasmids harboring candidate effector genes were introduced into competent *A. tumefaciens* strain GV3101 cells using the freeze-thaw transformation method. Transformed cells were selected on LB agar plates containing appropriate antibiotics and subsequently propagated in LB liquid medium at 28 °C with shaking at 180 rpm for 24 h to obtain sufficient biomass for infiltration assays.

Bacterial cultures were harvested by centrifugation (5000 rpm, 5 min) and washed three times with infiltration buffer composed of 10 mM MgCl_2_, 10 mM MES, and 100 μM acetosyringone (pH 5.7). The cell pellets were resuspended in the same buffer and adjusted to an optical density at an OD_600_ of 1.0, depending on the experimental requirements. Suspensions were incubated in the dark at 28 °C for 2-3 h to induce virulence gene expression prior to infiltration.

For transient expression assays, bacterial suspensions were syringe-infiltrated into the abaxial side of fully expanded leaves of 5-6-week-old *N. benthamiana* plants using a needleless syringe. Infiltrated plants were maintained under the same controlled growth conditions described above. Phenotypic responses, including HR development, tissue collapse, and other visible symptoms, were monitored at defined time points post-infiltration. All assays were performed with at least three biological replicates to ensure reproducibility.

### Yeast signal sequence trap system

The functionality of the predicted Ss1624 SP was evaluated using a yeast signal sequence trap assay as previously described (Yin *et al*., 2018). The pSUC2 vector, which contains a truncated invertase gene lacking both the initiation methionine and its native SP, was used as the reporter backbone. The predicted SP sequence of Ss1624 was fused in-frame upstream of the truncated invertase coding sequence, generating the construct pSUC2:Ss1624(SP)-Invertase.

As a positive control, the well-characterized SP from the *Phytophthora sojae* RXLR effector Avr1b was cloned into pSUC2 to produce pSUC2:Avr1b(SP)-Invertase. The empty pSUC2 vector lacking any SP and an untransformed yeast strain served as negative controls to validate assay specificity. Primers and constructs used for these analyses were listed in **Supplemental Tables 3 and 4**.

The constructs pSUC2:Ss1624(SP)-Invertase and pSUC2:Avr1b(SP)-Invertase were introduced into the yeast strain YTK12 using the lithium acetate transformation method. Transformants were initially selected on synthetic dropout medium lacking tryptophan (CMD-W). To assess invertase secretion, yeast colonies were subsequently transferred to YPRAA medium. Functional SPs enable secretion of invertase, which catalyzes the reduction of 2,3,5-triphenyltetrazolium chloride (TTC) to triphenylformazan, a red insoluble compound indicative of invertase enzymatic activity. The development of red coloration was used as a qualitative readout of SP functionality.

### VIGS in N. benthamiana

VIGS assays were conducted in *N. benthamiana* using a tobacco rattle virus (TRV)-based system as previously described (Velásquez *et al*., 2009). Constructs pTRV2-*GFP*, pTRV2-*NbBAK1*, and pTRV2-*NbSOBIR1* were generated and subsequently introduced into *A. tumefaciens* strain GV3101 via the freeze-thaw method.

*Agrobacterium* cultures carrying pTRV1 (RNA1) and pTRV2-derived constructs (RNA2) were grown separately in LB medium under antibiotic selection, then adjusted to an OD_600_ of 0.2. Equal volumes of pTRV1 and pTRV2 cultures were mixed in a 1:1 ratio prior to infiltration. The mixed suspensions were infiltrated into the two primary leaves of four-leaf-stage *N. benthamiana* seedlings using a needleless syringe.

Following infiltration, plants were maintained under controlled growth conditions for systemic gene silencing to occur. The gene silencing efficiency of *NbBAK1* and *NbSOBIR1* was evaluated by qRT-PCR with gene-specific primers, using samples collected 3 weeks after infiltration. Only plants exhibiting confirmed gene silencing were used for downstream functional assays to ensure the reliability of the phenotypic analyses. Experiments were repeated at least three times with independent biological replicates. Primers and constructs used for these analyses were listed in **Supplemental Tables 3 and 4**.

### RNA extraction and qRT-PCR analysis

Plant tissues or fungal cells were harvested at designated time points, immediately flash-frozen in liquid nitrogen, and ground into a fine powder using pre-chilled mortar and pestle. Total RNA was extracted using either the Qiagen RNeasy^®^ Mini Kit or Qiagen RNeasy^®^ Plant Mini Kit, following the manufacturer’s protocols. RNA quality was assessed by agarose gel electrophoresis to verify integrity, and RNA concentration was determined using a NanoDrop 2000 spectrophotometer (Thermo Fisher Scientific, USA).

First-strand cDNA synthesis was performed using 1 μg of total RNA with HiScript III RT SuperMix (Vazyme Biotech, Nanjing, China). Quantitative PCR reactions were carried out on an Applied Biosystems QuantStudio^TM^ 6 Flex Real-Time PCR System under the following thermal cycling conditions: an initial step at 50 °C for 2 min, followed by denaturation at 95 °C for 2 min; 40 amplification cycles consisting of 95 °C for 15 s and 60 °C for 30 s; and a final dissociation stage with 95 °C for 15 s, 60 °C for 1 min, and 95 °C for 15 s to verify amplification specificity.

Gene expression levels were normalized using the *S. scitamineum* β-actin gene and the *N. benthamiana* elongation factor 1-alpha (*EF1α*) gene as internal reference controls, depending on the biological system analyzed. Three independent biological replicates were included for each experimental condition. Relative transcript abundance was calculated using the 2^-ΔΔCt^ method. Statistical analyses were performed using unpaired Student’s *t* tests, and differences were considered statistically significant at *P* < 0.05. All primer sequences used in this study are provided in **Supplemental Table 3**.

### Co-immunoprecipitation and western blot analysis

HA-tagged or RFP-fused constructs were transiently expressed in *N. benthamiana* leaves via *A. tumefaciens*-mediated infiltration. Total protein extracts were prepared 48 h post-infiltration to allow sufficient accumulation of recombinant proteins. Leaf tissues were harvested, immediately frozen in liquid nitrogen, and ground to a fine powder using pre-chilled mortars and pestles. The resulting powder was homogenized in lysis buffer consisting of 150 mM NaCl, 25 mM Tris-HCl (pH 7.5), 10% glycerol, 5 mM dithiothreitol (DTT), 1 mM phenylmethylsulfonyl fluoride (PMSF), 0.5% NP-40, and 0.1% (v/v) protease inhibitor cocktail to minimize protein degradation.

Cell debris was removed by sequential centrifugation steps (13,000 rpm, 10 min per cycle, 4 °C), with the supernatant collected after each spin to ensure maximal recovery of soluble proteins. At least 500 μL of clarified protein extract was transferred into pre-chilled 1.5-mL tubes and incubated with 15 μL of RFP-Trap^®^ M magnetic beads (ChromoTek, Germany) at 4 °C for 1 h under gentle rotation to allow immunoprecipitation of RFP-tagged proteins and their interacting partners.

Following incubation, magnetic beads were retrieved using a magnetic separation rack and washed four times with 1 mL of pre-cooled wash buffer (150 mM NaCl, 25 mM Tris-HCl pH 7.5, 10% glycerol, 0.1% Tween 20, 5 mM DTT, 1 mM PMSF) to reduce nonspecific binding. Bound proteins were eluted by resuspending beads in 40 μL of phosphate-buffered saline (PBS, pH 7.4) and mixing with 10 μL of 2× SDS-PAGE loading buffer (120 mM Tris-HCl pH 6.8, 20% glycerol, 4% SDS, 0.04% bromophenol blue, 10% β-mercaptoethanol). Samples were denatured at 95 °C for 10 min prior to electrophoretic separation.

Proteins were resolved on 10% or 12% SDS-polyacrylamide gels at a constant voltage of 80 V for 90-120 min, depending on protein size. Separated proteins were transferred onto methanol-activated polyvinylidene fluoride (PVDF) membranes using a wet/tank transfer system (Bio-Rad) according to the manufacturer’s instructions. Membranes were blocked in Tris-buffered saline (TBS) containing 0.1% Tween 20 and 5% (w/v) skim milk for 1 h at room temperature to prevent nonspecific antibody binding.

Subsequently, membranes were incubated with primary antibodies, including mouse monoclonal anti-RFP antibody (1:5000 dilution; ChromoTek, Germany) or mouse monoclonal anti-HA antibody (1:5000 dilution; Abmart, China), overnight at 4 °C. After extensive washing, membranes were incubated with horseradish peroxidase (HRP)-conjugated goat anti-mouse IgG secondary antibody (1:10,000 dilution; Abmart, China) for 1 h at room temperature. Immunoreactive bands were visualized using Clarity^TM^ Western ECL substrate (Bio-Rad, Hercules, CA, USA) and detected using a chemiluminescence imaging system following the manufacturer’s instructions. Band intensities were interpreted as indicators of protein accumulation and interaction, and representative blots were selected from at least two independent experiments.

### Pathogenicity assays

Sugarcane smut infection. WT *S. scitamineum* strains (*MAT-1* and *MAT-2*) and their derivative mutants were cultured on YEPS agar plates for 48 h, followed by liquid YEPS culture at 28 °C with shaking at 180 rpm for 36 h to obtain actively growing cells. Cultures were harvested by centrifugation, washed with sterile water, and resuspended to an optical density at 600 nm (OD_600_) of 1.0. Compatible haploid strains (*MAT-1* and *MAT-2*) were mixed in equal proportions immediately prior to inoculation to ensure mating compatibility (Cai *et al*., 2021). Sugarcane cultivar ROC22 was used as the host plant. Seedlings were pre-germinated and grown in a greenhouse at 28 ± 2 °C under a 14 h light/10 h dark photoperiod until reaching the 3-4 leaf stage. Inoculation was performed using a needle puncture method, in which the stem of each seedling was gently pierced with a sterile 1-mL syringe, followed by injection of 200 μL of fungal suspension into the stem parenchyma. Each treatment included at least 30 plants, and the entire experiment was independently repeated three times to ensure reproducibility. Disease development was monitored at defined intervals, and phenotypic progression was recorded systematically.

Rice bacterial blight assay. Pathogenicity assays for bacterial blight were conducted using the leaf-clipping method (Ma & Yuan, 2018). An overnight culture of *Xoo* was centrifuged, and the bacterial pellet was resuspended in 10 mM MgCl_2_ to an OD_600_ of 0.8. Leaf tips of 6-week-old rice plants were clipped with scissors dipped in the bacterial suspension. Inoculated plants were maintained under standard growth conditions, and lesion lengths were measured at 14 dpi as a quantitative indicator of disease severity.

Rice blast assay. For rice blast infection, *M. oryzae* was cultured on oatmeal agar plates at 28 °C in the dark (Cai *et al*., 2025). Conidia were harvested by flooding plates with sterile water and filtering through Miracloth to remove mycelial debris. Four-leaf-stage rice seedlings were spray-inoculated with a conidial suspension adjusted to 1 × 10^6^ conidia/mL in 0.1% (v/v) Tween 20. Following inoculation, plants were incubated in a growth chamber at 26 °C under high humidity conditions with a 12 h light/12 h dark photoperiod. Disease symptoms were documented at 7 dpi, and lesion development was used to compare susceptibility among genotypes.

Rice sheath blight assay (Ni *et al*., 2026). *R. solani* was cultured on PDA at 28 °C in darkness for 2–3 days. Mycelial plugs (5 mm diameter) were transferred to fresh PDA plates pre-laid with sterile filter paper strips (1× 0.2 cm) and incubated under the same conditions for 3 days until the strips were fully colonized. These colonized strips were then used as inoculum and embedded into rice sheaths at the late tillering stage. Disease severity was assessed 7 dpi.

### Fungal growth and stress sensitivity assays

WT *S. scitamineum* strains (*MAT-1* and *MAT-2*) and their derivative mutants were evaluated for growth under a range of environmental stresses. Cultures were grown on YEPS medium supplemented with stress-inducing agents, including 250 mM CaCl_2_, 0.5 mM H_2_O_2_, 10 mM FeSO_4_, 10 mg/mL Congo Red, 1.5 mM CuSO_4_, 0.002% SDS, 0.5 M KCl, 1 M sorbitol, or 0.5 M NaCl. These conditions were selected to assess tolerance to oxidative stress, osmotic stress, membrane perturbation, and cell wall integrity challenges.

In parallel, fungal growth was also evaluated on minimal medium (MM) to assess baseline growth capacity. Plates were incubated at 28 °C for 3 days, after which colony morphology, pigmentation, and radial growth rates were recorded. Observations were captured using a Leica M165 FC stereomicroscope to ensure consistent documentation of morphological differences. Growth measurements were compared across strains, and each assay was performed at least twice independently, with three biological replicates per condition.

### Split luciferase complementation (SLC) assay

Ss1624 and RLK15 coding sequences lacking their SPs were cloned into pCAMBIA1300-based vectors carrying the N-or C-terminal luciferase fragments to generate Ss1624-nLuc and cLuc-RLK15. Primers and constructs used for these analyses were listed in **Supplemental Tables 3 and 4**.

The constructs were introduced into *A. tumefaciens* strain GV3101 and co-expressed in *N. benthamiana* leaves via agroinfiltration, following procedures described above. Three days after infiltration, infiltrated leaf regions were uniformly sprayed with 0.3 mg/mL D-luciferin sodium salt (Coolaber, China, CL6929). Leaves were then incubated in the dark for 3 min to allow substrate penetration and stabilization of the luminescent reaction.

Luciferase activity was detected using a low-light cooled charge-coupled device (CCD) imaging system (NightSHADE LB 985; Berthold Technologies, Germany). Luminescence was used as a readout of split-luciferase complementation. Representative images were obtained from independent biological replicates.

### Analysis of Ss1624 and RLK15 gene expression patterns

To examine temporal expression patterns of *Ss1624* and *RLK15* during infection, sugarcane stem tissues were sampled following the same pathogenicity assay framework described above. Briefly, tissues were collected at 2, 4, 6, 8, 10, 12, 14, and 18 dpi from plants inoculated with WT, Δ*ss1624*, and *Ss1624 OE* strains. Sampling at multiple time points enabled the capture of dynamic transcriptional changes associated with early colonization, progression of infection, and later-stage host responses.

Total RNA was extracted from each sample using standard plant RNA isolation procedures, followed by DNase treatment to remove residual genomic DNA. First-strand cDNA synthesis was performed using reverse transcriptase according to the manufacturer’s instructions. qRT-PCR was then conducted to determine relative transcript abundance of *Ss1624* or *RLK15*. Gene-specific primers were used, and amplification efficiency was verified prior to analysis (**Supplemental Table 3**). Expression levels were normalized using the *S. scitamineum* β-actin gene, and relative expression was calculated using the 2^-ΔΔCt^ method. Each biological sample was analyzed in technical triplicate to ensure reproducibility, and at least three independent biological replicates were included per treatment.

### ROS and callose staining

To assess host immune responses, ROS accumulation and callose deposition were evaluated in *N. benthamiana* leaves. For ROS detection, leaves were harvested at 48 hpi and incubated in 1 mg/mL 3,3′-diaminobenzidine (DAB) solution in the dark for 12-16 h. DAB polymerizes in the presence of hydrogen peroxide, forming a brown precipitate indicative of ROS accumulation. After staining, chlorophyll was removed by boiling the tissues in anhydrous ethanol for 10 min, allowing clearer visualization of staining patterns.

For callose deposition analysis, leaf segments (approximately 1 cm × 2 cm) were excised and fixed in Carnoy’s solution (anhydrous ethanol: acetic acid: trichloromethane, 6:3:1, v/v/v) for 24 h to preserve cellular structure and polysaccharide deposits. Samples were subsequently destained by heating in anhydrous ethanol for 10 min to remove pigments. Callose was then stained with 0.01% (w/v) aniline blue prepared in 150 mM K_2_HPO_4_ buffer and incubated in the dark for 6-8 h. Callose deposits were visualized under ultraviolet excitation using a fluorescence microscope, and representative images were captured under identical exposure settings for comparative analysis. The number of callose deposits was quantified in a single microscopic field per sample, with at least three random fields examined for each biological replicate. Data are expressed as mean ± SD and analyzed statistically.

### Phosphorylation assay

Kinase activity assays were performed following the protocol described by Sung *et al*. with minor modifications (Sung *et al*., 2025). Reactions were assembled in kinase buffer containing 20 mM Tris-HCl (pH 7.5), 10 mM MgCl_2_, 1 mM CaCl_2_, 1 mM dithiothreitol, and 100 μM ATP. Recombinant Ss1624 protein (2 μg per reaction) was incubated with recombinant RLK15 ^C^-terminal protein at varying concentrations (0, 1.2, 7.2, and 12.2 μg) to assess dose-dependent phosphorylation.

Reactions were incubated at 30 °C for 30 min to allow kinase activity. Subsequently, samples were denatured and resolved by SDS-PAGE under standard conditions. Proteins were transferred to PVDF membranes and probed using an anti-phospho-(Ser/Thr) antibody (Abcam, ab117253) to detect phosphorylation events. Signals were visualized using appropriate chemiluminescent detection reagents, and phosphorylation levels were compared across different substrate concentrations to evaluate the kinase-dependent modification of RLK15 ^C^ under the tested in vitro conditions. Each assay was performed in at least three independent experiments to confirm consistency.

### In vitro protein expression and purification

The *Ss1624* coding sequence was cloned into the pPIC9K-His expression vector for secretion in *P. pastoris*. The construct was linearized and transformed into *P. pastoris* strain GS115 competent cells via electroporation. Transformants were selected on minimal dextrose (MD) plates, and clones carrying multiple insertions were further screened using increasing concentrations of G418 to identify high-copy integrants.

Positive clones were cultured in buffered methanol-complex medium (BMMY), and protein expression was induced with methanol supplementation over 96 h. The culture supernatant containing secreted His-tagged Ss1624 protein was collected by centrifugation and clarified through filtration. Purification was performed using nickel-nitrilotriacetic acid (Ni-NTA) affinity chromatography under native conditions. Eluted fractions containing the target protein were pooled, concentrated, and buffer exchanged into phosphate-buffered saline (PBS) using a 10 kDa molecular weight cutoff centrifugal filter. Purified protein aliquots were stored at -80 °C for subsequent analyses.

For RLK15 C-terminal protein expression, the corresponding coding region was cloned into the pET28a-MBP-His vector and expressed in *E. coli* strain (*Trans*etta DE3). Expression was induced with 0.2 mM IPTG at 16 °C for 16 h to promote soluble protein production. Cells were harvested by centrifugation and lysed by sonication in lysis buffer. The clarified lysate was incubated with Ni-NTA resin for affinity purification. After extensive washing to remove nonspecifically bound proteins, the target fusion protein was eluted, concentrated, and further purified. The final protein was exchanged into PBS and stored in aliquots at -80 °C to maintain stability for downstream biochemical assays.

### Generation of transgenic plants

All transgenic rice lines used in this study were generated in the *japonica* cultivar Zhonghua 11 (ZH11) background. CRISPR–Cas9-mediated mutagenesis of OsRLK15 was performed using gene-editing constructs that were designed and transformed into ZH11 by Biogle Gene Technology. Mutant candidates were identified by PCR amplification and Sanger sequencing of the target regions from leaf-extracted genomic DNA. For overexpression, the full-length coding sequence of OsRLK15 was inserted in frame with GFP into the pCAMBIA super1300 vector via SmaI and KpnI restriction sites. *Agrobacterium tumefaciens* EH105 mediated transformation was carried out by BioRun Biotechnology. Positive transgenic plants were confirmed by immunoblotting or semiquantitative RT-PCR. Primers and constructs are listed in **Supplemental Tables 3 and 4**.

### Phylogenetic analyses

For phylogenetic analysis of plant RLK15, protein sequences of all CrRLK1L-1 type receptor-like kinases were retrieved from the metaRLK database. Initial sequence alignment was performed using MUSCLE v5.2 with the -super5 option. The alignment was subsequently trimmed using trimAl v1.4 with the -gappyout parameter. A preliminary phylogenetic tree was constructed using VeryFastTree v4.0.5 to identify and extract the well-supported RLK15 clade containing both monocot and eudicot homologs. For high-resolution phylogenetic reconstruction of this specific RLK15 subfamily, the extracted sequences were re-aligned using MUSCLE v5.2 (using the-align option), trimmed with trimAl v1.4 (-gappyout), and the final maximum-likelihood phylogenetic tree was inferred using IQ-TREE 3.

For phylogenetic analysis of the fungal effector Ss1624, homologous protein sequences corresponding to the fungi-level orthogroup ‘400260at4751’ were retrieved from the OrthoDB v12.2 database. These protein sequences were aligned using MUSCLE v5.2 (-align option) and trimmed with trimAl v1.4 (-gappyout). The phylogenetic tree for the Ss1624 orthogroup was subsequently reconstructed using IQ-TREE 3. Sequence identity between Ss1624 and its homologs was calculated based on the generated multiple sequence alignments.

### Statistical analysis

Data were analyzed using GraphPad Prism version 9.5 (GraphPad Software, San Diego, CA, USA). Comparisons between two groups were performed using Student’s *t* test. Multiple-group comparisons were analyzed by analysis of variance (ANOVA) followed by Tukey’s honestly significant difference (HSD) test. Differences were considered statistically significant at P < 0.05. Data are presented as means ± standard deviations (SD) from at least three independent replicates, unless otherwise indicated.

## Data Availability Statement

All data needed to evaluate the conclusions in the paper are present in the paper and/or the Supplementary Materials.

## Supplementary Data

**Supplemental Figure 1.**
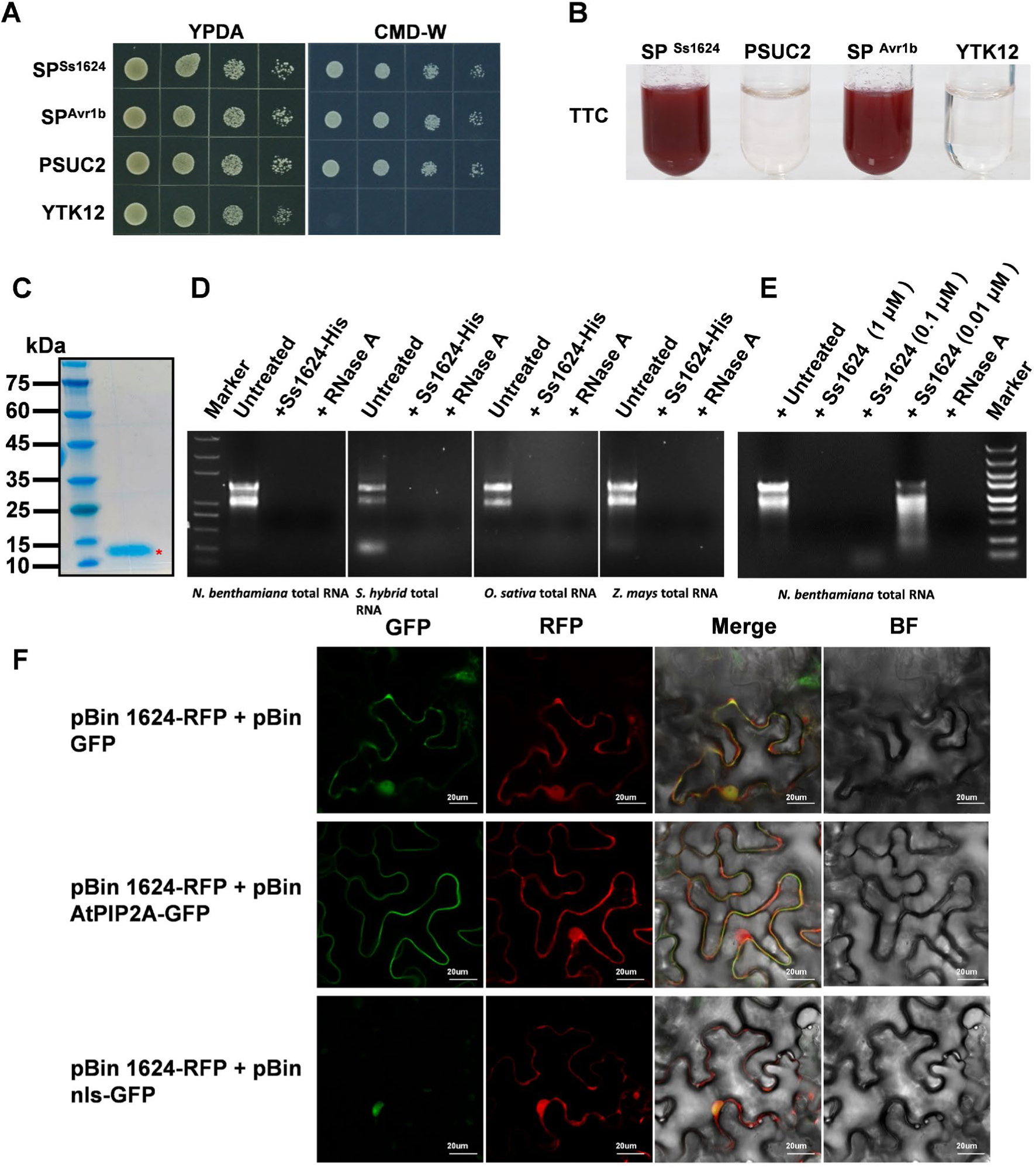
Ss1624 from *S. scitamineum* encodes a ribonuclease that induces cell death. (**A**) Yeast signal sequence trap assay assessing secretion activity. The predicted SP of Ss1624 was fused to a secretion reporter and expressed in yeast strain YTK12. Avr1b SP served as a positive control, whereas wild-type (WT) YTK12 and empty vector transformants served as negative controls. Growth on CMD-W medium indicates functional secretion. (**B**) Invertase secretion was further confirmed using the 2,3,5-triphenyltetrazolium chloride (TTC) assay. Fusion of Ss1624 or Avr1b SPs to yeast invertase enables extracellular secretion and reduction of TTC to red 1,3,5-triphenylformazan. (**C**) Expression and purification of recombinant Ss1624-His protein in *Pichia pastoris*. SDS-PAGE analysis stained with Coomassie brilliant blue showed successful expression and secretion of recombinant Ss1624-His protein (predicted molecular weight is approximately 14.25 kDa). Protein accumulation in the culture medium was highest at 72-96 h following methanol induction. The target protein was subsequently purified using Ni-NTA affinity chromatography, yielding a single major band corresponding to the expected molecular weight after purification. (**D**) Recombinant Ss1624 exhibits ribonuclease activity *in vitro*. Purified Ss1624 protein expressed in *P. pastoris* was incubated with total RNA extracted from *N. benthamiana*, sugarcane (*Saccharum* spp. hybrid), rice (*O. sativa*), and maize (*Z. mays*), alongside RNase A as a positive control and untreated RNA as a negative control. (**E**) Dose-dependent RNA degradation by Ss1624. Total RNA from *N. benthamiana* was incubated with increasing concentrations of Ss1624 protein (1, 0.1, and 0.01 μM), showing detectable degradation at all tested concentrations, with 0.01 μM representing the lowest effective level. (**F**) Subcellular localization of Ss1624-RFP was examined by co-expression with AtPIP2A (a plasma membrane marker), nls-GFP (a nuclear marker), or GFP in *N. benthamiana* leaf cells. Fluorescence signals were observed using confocal microscopy at 36 h post-infiltration (hpi). For each panel set (left to right), images show the GFP channel, RFP channel, merged fluorescence, and corresponding bright-field (BF) image. Co-localization is indicated by yellow fluorescence in the merged images. Scale bars = 20 μm.

**Supplemental Figure 2.**
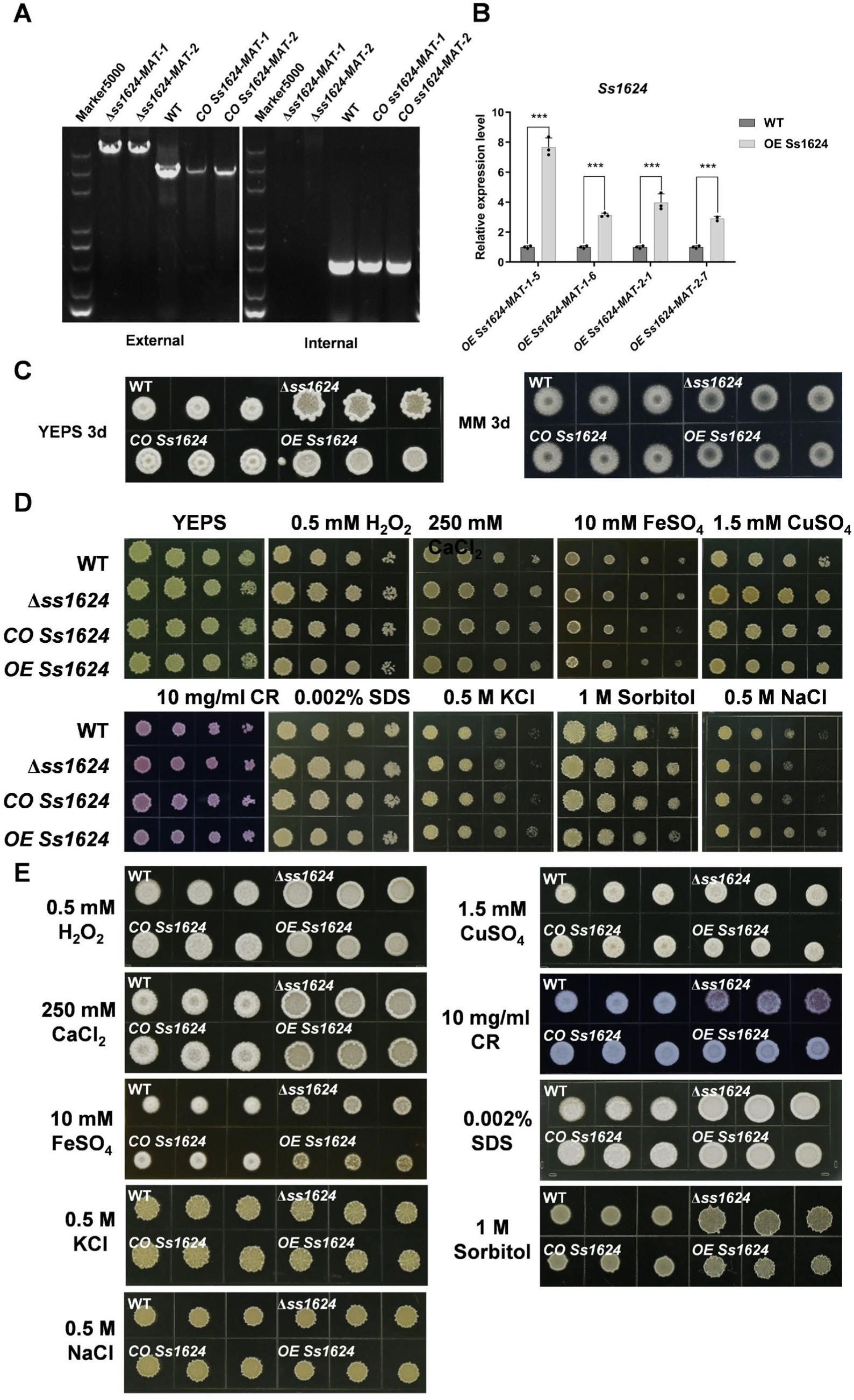
Phenotypic analysis of *Ss1624* knockout, complementation, and overexpression strains under nutritional and stress conditions. (**A**) PCR-based verification of *Ss1624* gene manipulation. Genomic DNA from wild-type (WT), two independent knockout mutants (Δ*ss1624-MAT-1* and Δ*ss1624-MAT-2*), and complemented strains (*CO Ss1624-MAT-1* and *CO Ss1624-MAT-2*) was amplified using external and internal primer sets. External primers flank the *Ss1624* locus and generate amplicons only when the native gene is present or restored, whereas internal primers specifically amplify the *Ss1624* coding sequence, producing bands only in strains harboring the gene. The resulting band patterns confirmed successful gene deletion and complementation. (**B**) Relative expression of *Ss1624* determined by qRT-PCR. Transcript levels were significantly elevated in overexpression strains (*OE Ss1624-MAT-1* and *OE Ss1624-MAT-2*) compared with the wild type. Data are presented as mean ± SD from three biological replicates, with statistical significance assessed using Student’s *t* test (***, *p* < 0.001). (**C**) Growth of *Ss1624* mutant strains on different media. Colony morphology of WT, Δ*ss1624*, *CO Ss1624*, and *OE Ss1624* strains was examined after 3 days of incubation at 28 °C on YEPS and minimal medium (MM) plates. (**D** and **E**) Stress sensitivity assays of haploid and hyphal growth on YEPS agar supplemented with H_2_O_2_, CaCl_2_, FeSO_4_, CuSO_4_, Congo red, sorbitol, NaCl, KCl, and SDS. Plates were incubated at 28 °C in the dark for 3 days prior to imaging.

**Supplemental Figure 3.**
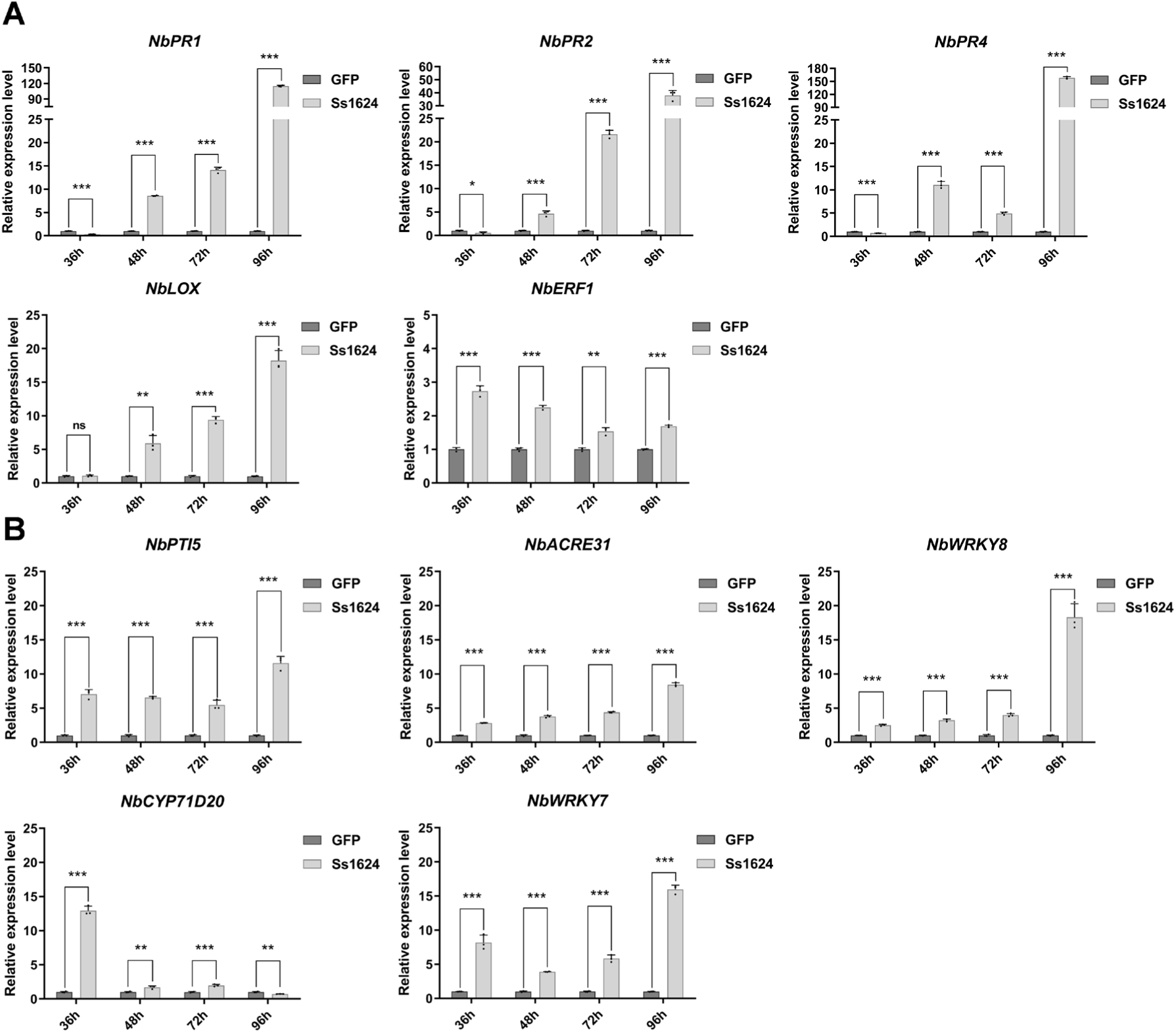
Ss1624 triggers plant immune responses in *N. benthamiana*. (**A**) Expression of defense-related marker genes (*NbPR1*, *NbPR2, NbPR4*, *NbLOX*, and *NbERF1*) was analyzed by qRT-PCR at the indicated time points. (**B**) Expression of PAMP-triggered immunity (PTI) marker genes (*NbPti5*, *NbACRE31*, *NbWRKY7*, *NbWRKY8*, and *NbCYP71D20*) was assessed by qRT-PCR. For all qRT-PCR analyses, data represent mean ± SD (n = 3). Statistical significance was determined by multiple unpaired t-tests with Benjamini, Krieger and Yekutieli (BKY) method to control the false discovery rate (FDR). Statistical significance between Ss1624 and GFP at each time point is indicated (\**, p* < 0.05; *\*\*, p* < 0.01; *\*\*\*, p* < 0.001; ns, not significant; Student’s *t* test).

**Supplemental Figure 4.**
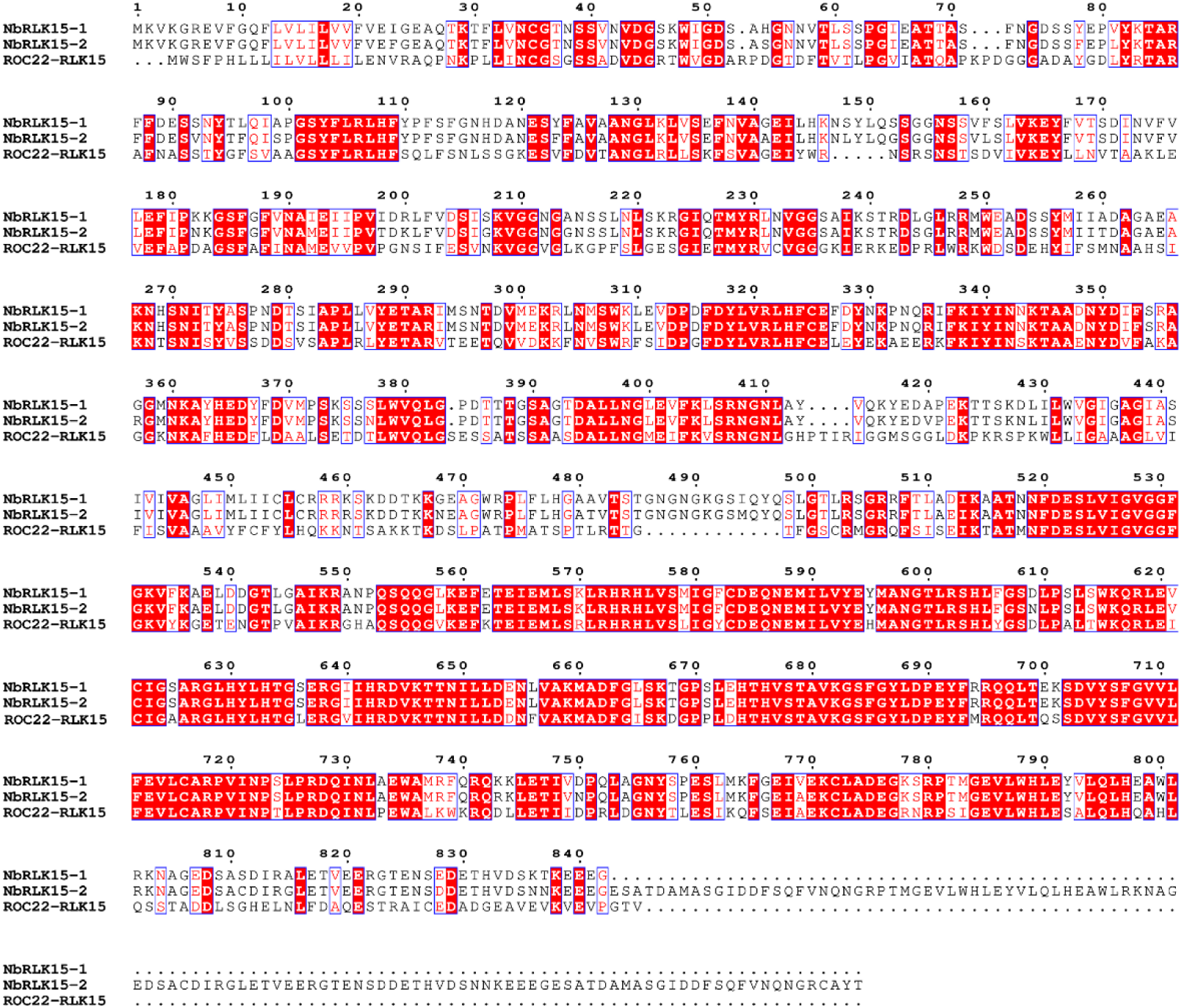
Sequence alignment of ROC22 sugarcane RLK15 with its two closest homologs in *N. benthamiana* (*NbRLK15-1* and *NbRLK15-2*). Sequence alignments were generated using MEGA11 and visualized with ESPript 3.2, with identical and highly conserved residues highlighted in color.

**Supplemental Figure 5.**
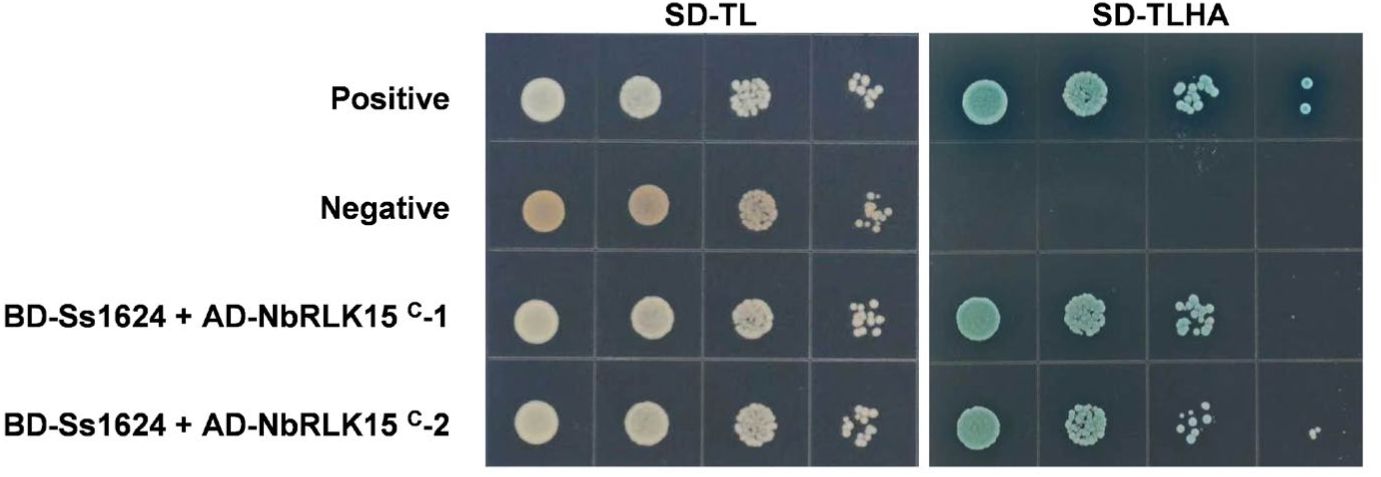
Yeast two-hybrid analysis showing that Ss1624 interacts with the intracellular kinase domain (C) of the *N. benthamiana* receptor kinase NbRLK15. Yeast co-transformants expressing BD-Ss1624 together with either AD-NbRLK15 ^C^-1 or AD-NbRLK15 ^C^-2 exhibited robust growth on both selective medium (SD/-T/-L/-H/-A) and non-selective medium (SD/-T/-L). The positive control (BD-53 + AD-T) showed strong growth on selective medium, whereas the negative control (BD-Lam + AD-T) failed to grow, confirming the specificity of the interaction.

**Supplemental Figure 6.**
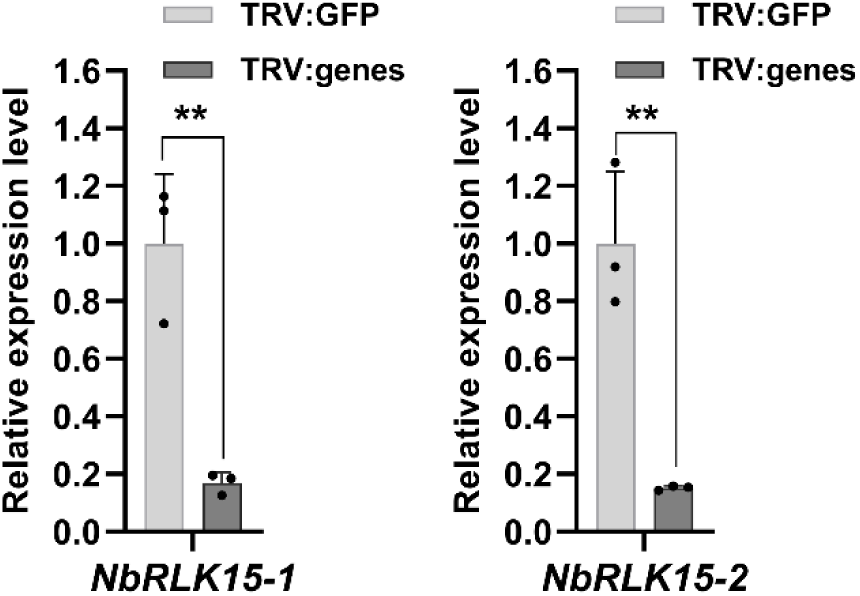
Validation of *NbRLK15* homologs silencing efficiency by qRT-PCR. Relative transcript levels of *NbRLK15-1* and *NbRLK15-2* were measured in *N. benthamiana* plants subjected to VIGS. The transcript abundance was significantly reduced in TRV:*genes* (TRV:*NbRLK15-1* or TRV:*NbRLK15-2*) compared to the TRV:*GFP* control. Data are presented as mean ± SD (n = 3), with statistical significance indicated (**, *p* < 0.01; Student’s *t* test).

**Supplemental Figure 7.**
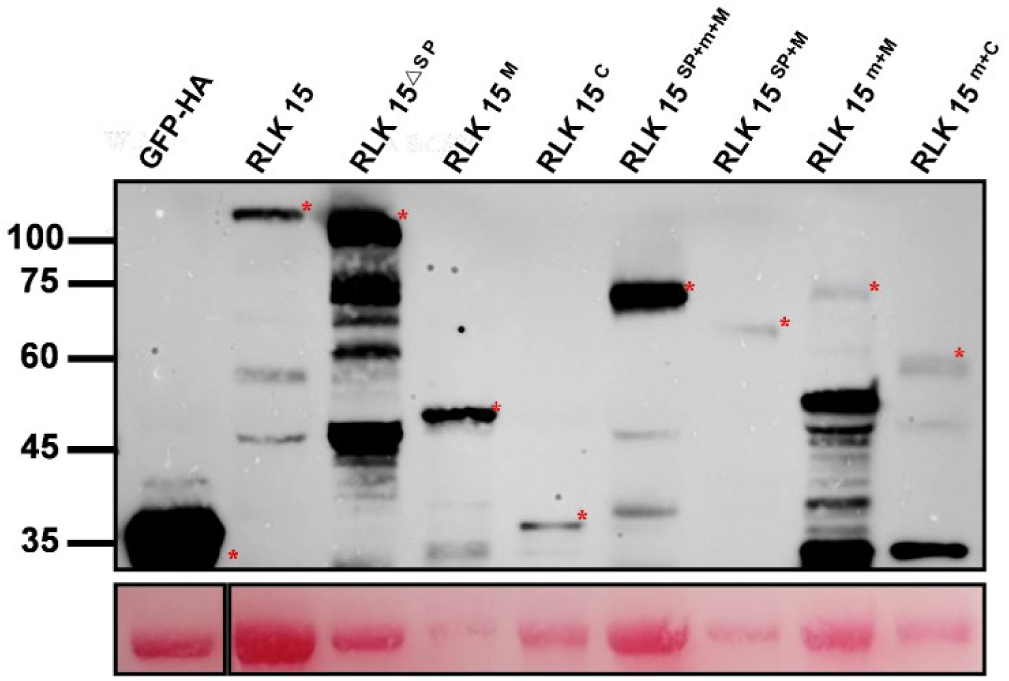
Western blot analysis confirming the expression of RLK15 constructs in *N. benthamiana* leaves. *Agrobacterium*-mediated transient expression was used to express the indicated RLK15 constructs in *N. benthamiana* leaves. Total protein extracts from infiltrated tissues were analyzed by immunoblotting using an anti-HA antibody. Ponceau S staining of the membrane was used as a loading control to verify equal protein loading.

**Supplemental Figure 8.**
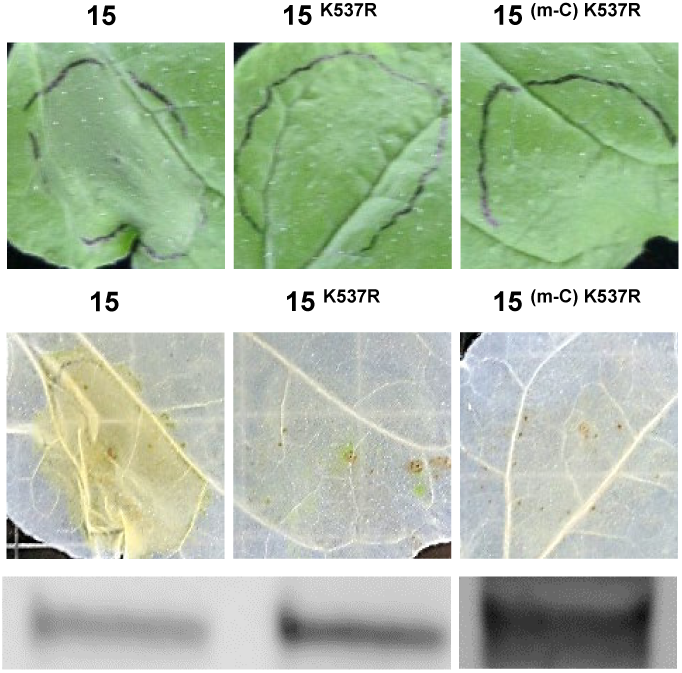
The kinase activity of RLK15 is required to trigger cell death. Cell death assay demonstrates that the K537R kinase-dead mutation abolishes the ability of RLK15 to induce cell death.

**Supplemental Figure 9.**
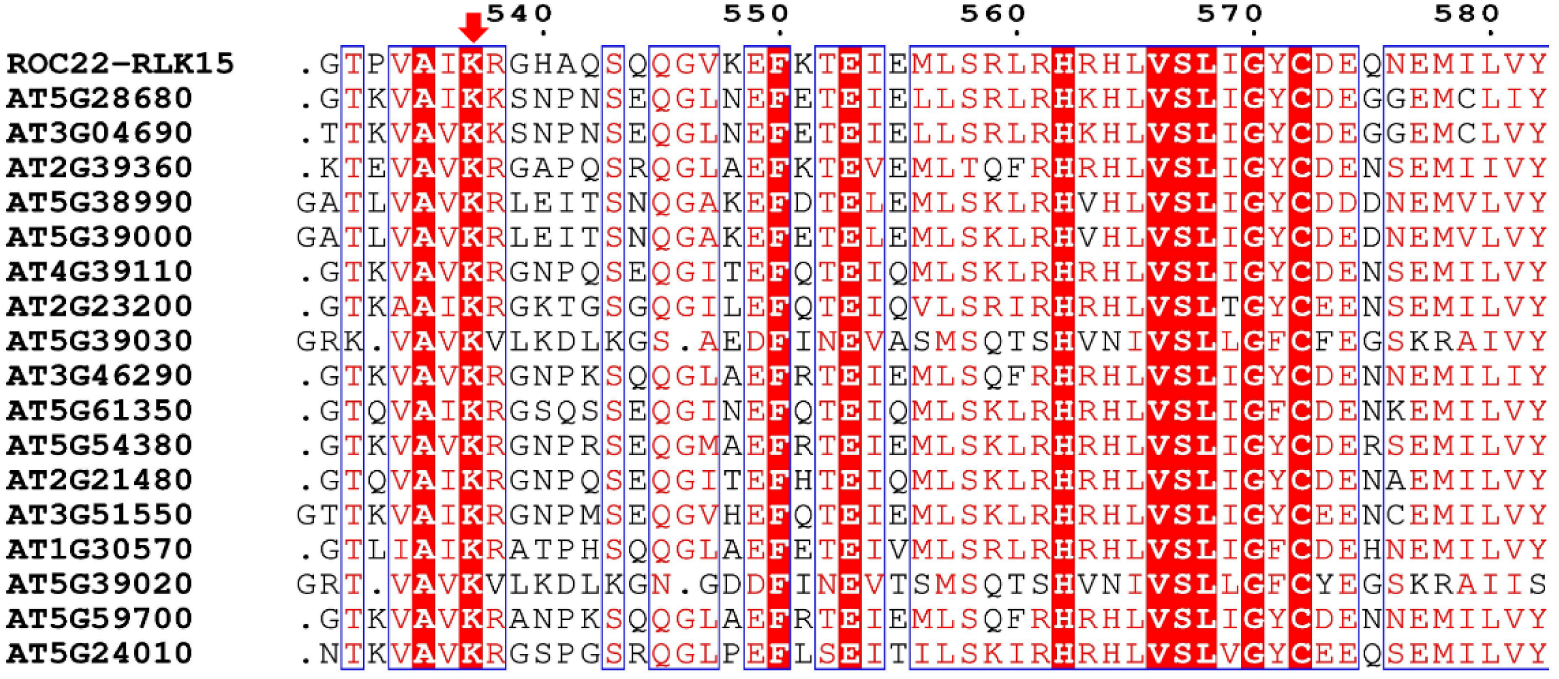
Multiple sequence alignment of RLK15 with members of the *Arabidopsis thaliana* CrRLK1L family. The alignment highlights a conserved region (amino acid positions 531-582) encompassing residues associated with kinase activity. Protein sequences were retrieved from The Arabidopsis Information Resource (TAIR). Alignments were generated using MEGA12 and visualized with ESPript 3.2, with identical and highly conserved residues indicated in color.

**Supplemental Figure 10.**
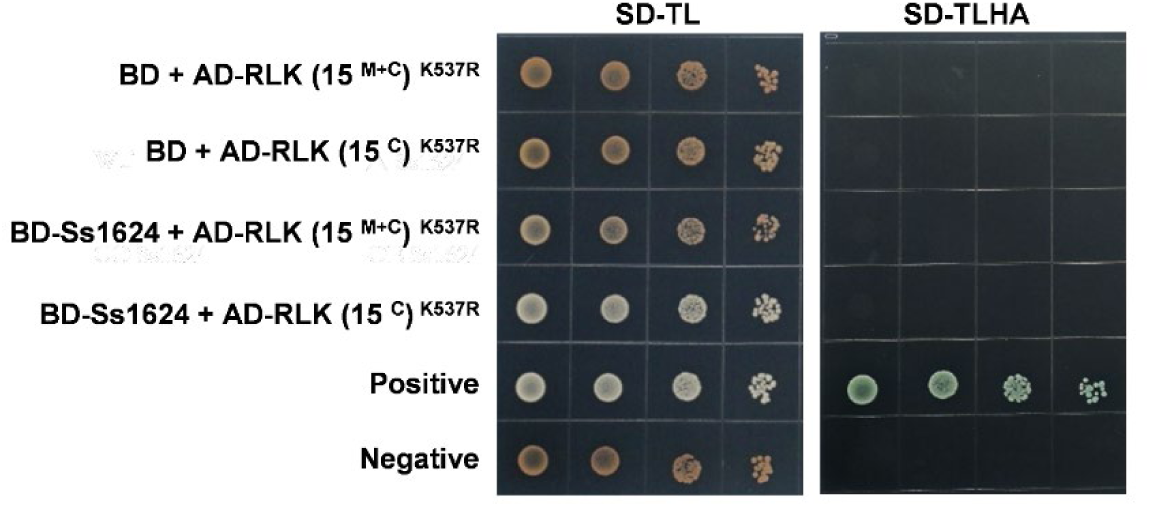
Yeast two-hybrid analysis showing that Ss1624 does not interact with kinase-dead RLK15 mutants. Yeast co-transformants expressing BD-Ss1624 together with AD-RLK15 ^(M+C)^ ^K537R^ or AD-RLK15 ^(C)^ ^K537R^ failed to grow on selective medium (SD/-T/-L/-H/-A), whereas normal growth was observed on non-selective medium (SD/-T/-L). The positive control (BD-53 + AD-T) showed robust growth on selective medium, while the negative control (BD-Lam + AD-T) did not grow, consistent with expected interactions.

**Supplemental Figure 11.**
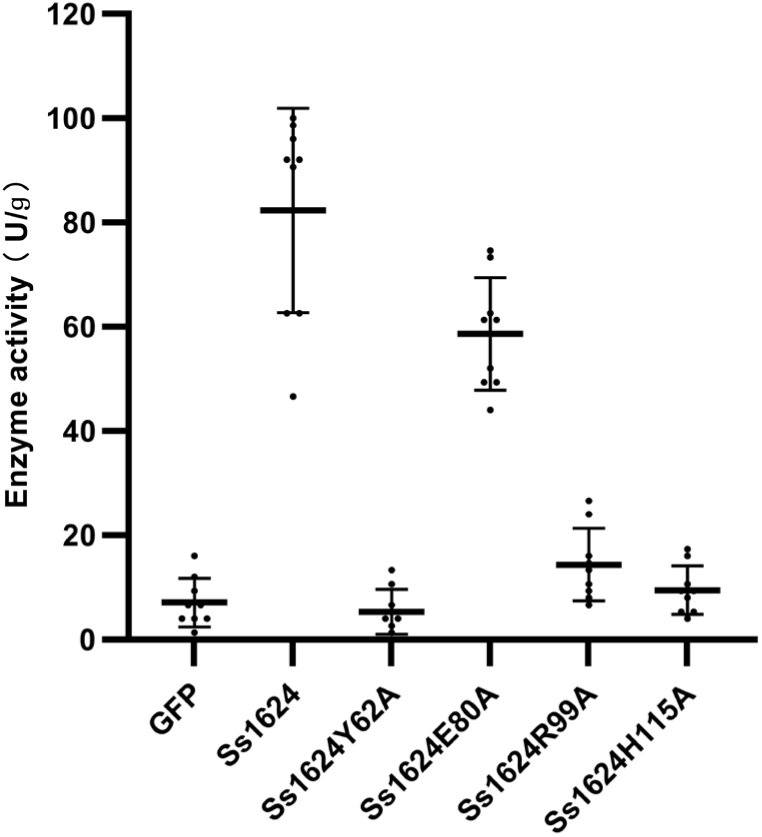
Comparative analysis of RNase activity between Ss1624 and its point mutants at key catalytic residues. Enzyme activity was assayed in sodium acetate buffer (pH 5.0) at 50 °C for 30 min, using yeast RNA at a final concentration of 0.15 mg/mL as substrate, and determined according to the national standard GB/T 34222-2017 (U/g). Data are presented as box plots (n ≥ 6).

**Supplemental Figure 12.**
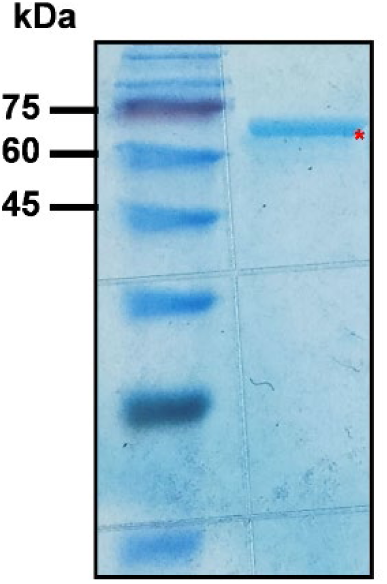
Prokaryotic expression and purification of recombinant RLK15 _C_. The coding sequence of RLK15 ^C^ was cloned into the pET28a-MBP vector with an N-terminal His tag and expressed in *E. coli* strain (*Trans*etta DE3). Following cell lysis, the soluble fraction was subjected to Ni-NTA affinity chromatography, including binding, washing, and elution steps. SDS-PAGE analysis confirmed successful expression and purification, with a prominent band at the expected molecular weight of 70.8 kDa observed in the eluate.

**Supplemental Figure 13.**
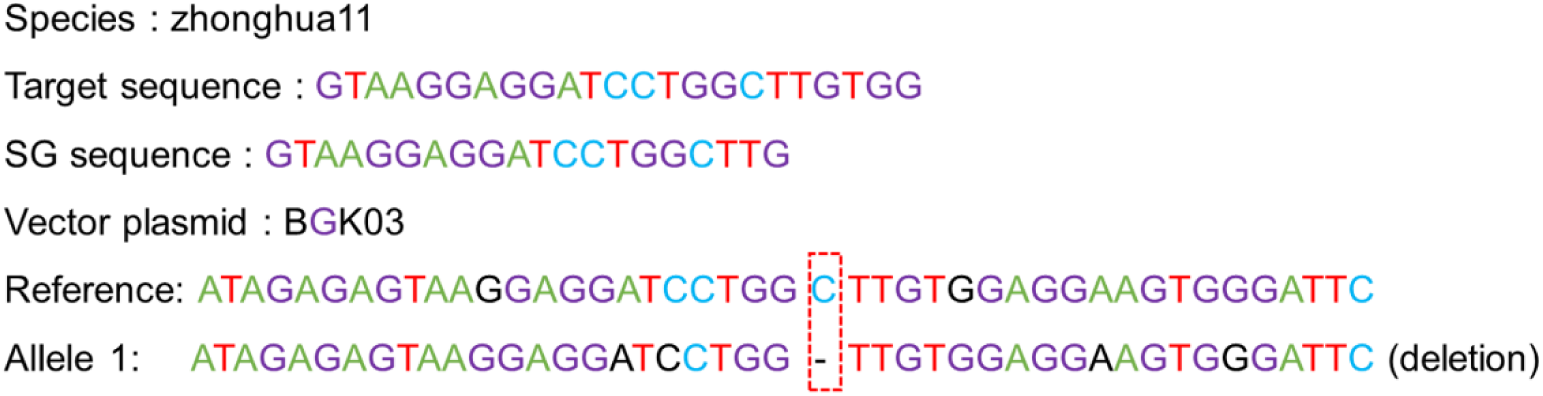
Schematic representation of CRISPR/Cas9-mediated mutations in the OsRLK15 locus of rice cultivar Zhonghua 11. The target sequence (GTAAGGAGGATCCTGGCTTGTGG) and corresponding sgRNA (GTAAGGAGGATCCTGGCTTG) within the BGK03 construct are shown. Sequencing was performed on the edited T2 plants, and the result was identified as a homozygous 1-bp deletion of a cytosine.

**Supplemental Figure 14.**
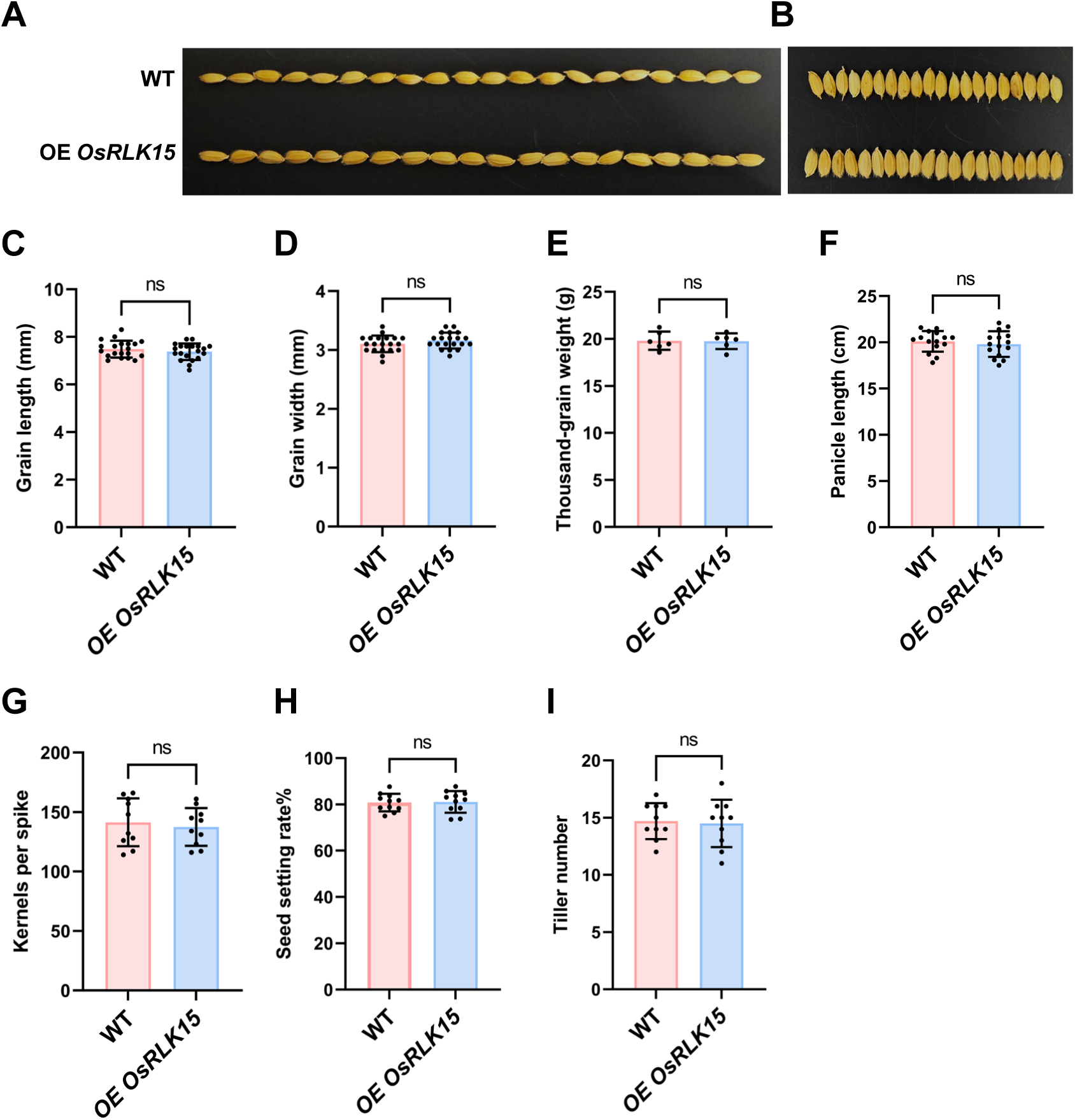
Effects of OsRLK15 overexpression on grain morphology and yield-related traits. (**A** and **B**) Grain morphology in WT and *OE OsRLK15* plants. Representative images illustrate grain length (A) and width (B) measured from 20 grains per line. (**C** and **D**) Quantitative analysis of grain length (C) and grain width (D). Data are mean ± SD (n = 20). (**E**) Thousand-grain weight. Data are mean ± SD (n = 6). (**F**) Panicle length. Data are mean ± SD (n = 15). (**G**) Number of kernels per spike. Data are mean ± SD (n = 10). (**H**) Seed setting rate. Data are mean ± SD (n = 11). (**I**) Tiller number. Data are mean ± SD (n = 10). Statistical comparisons with WT are indicated (ns, not significant; Student’s *t* test).

**Supplemental Figure 15.**
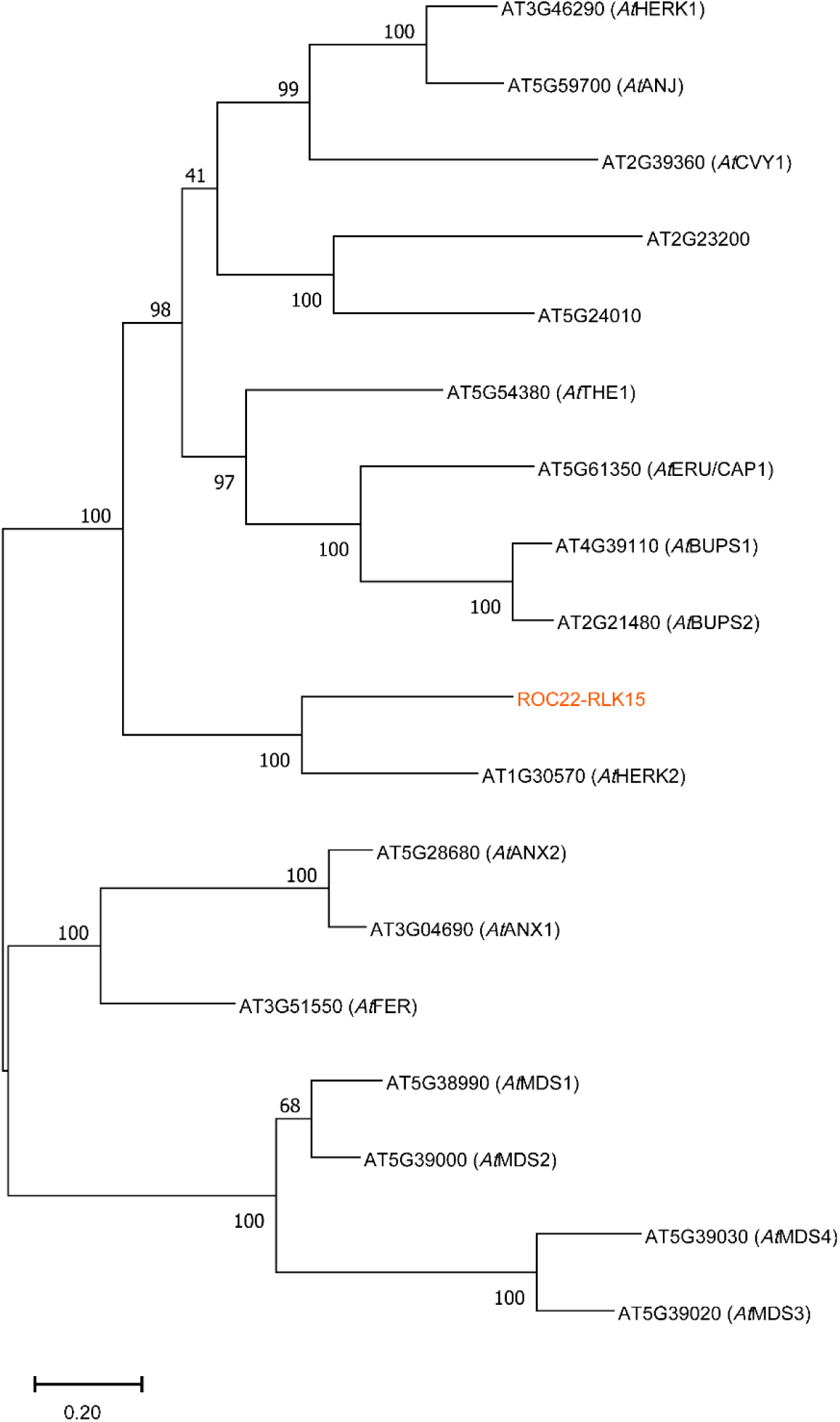
Phylogenetic analysis of *A. thaliana* CrRLK1L family members with the candidate protein ROC22-RLK15. The tree was constructed using the Maximum Likelihood (ML) method based on protein sequence alignments generated by MEGA11. Bootstrap values (≥40%) are indicated at the internal nodes. The scale bar represents 0.20 amino acid substitutions per site.

**Supplemental Figure 16.**
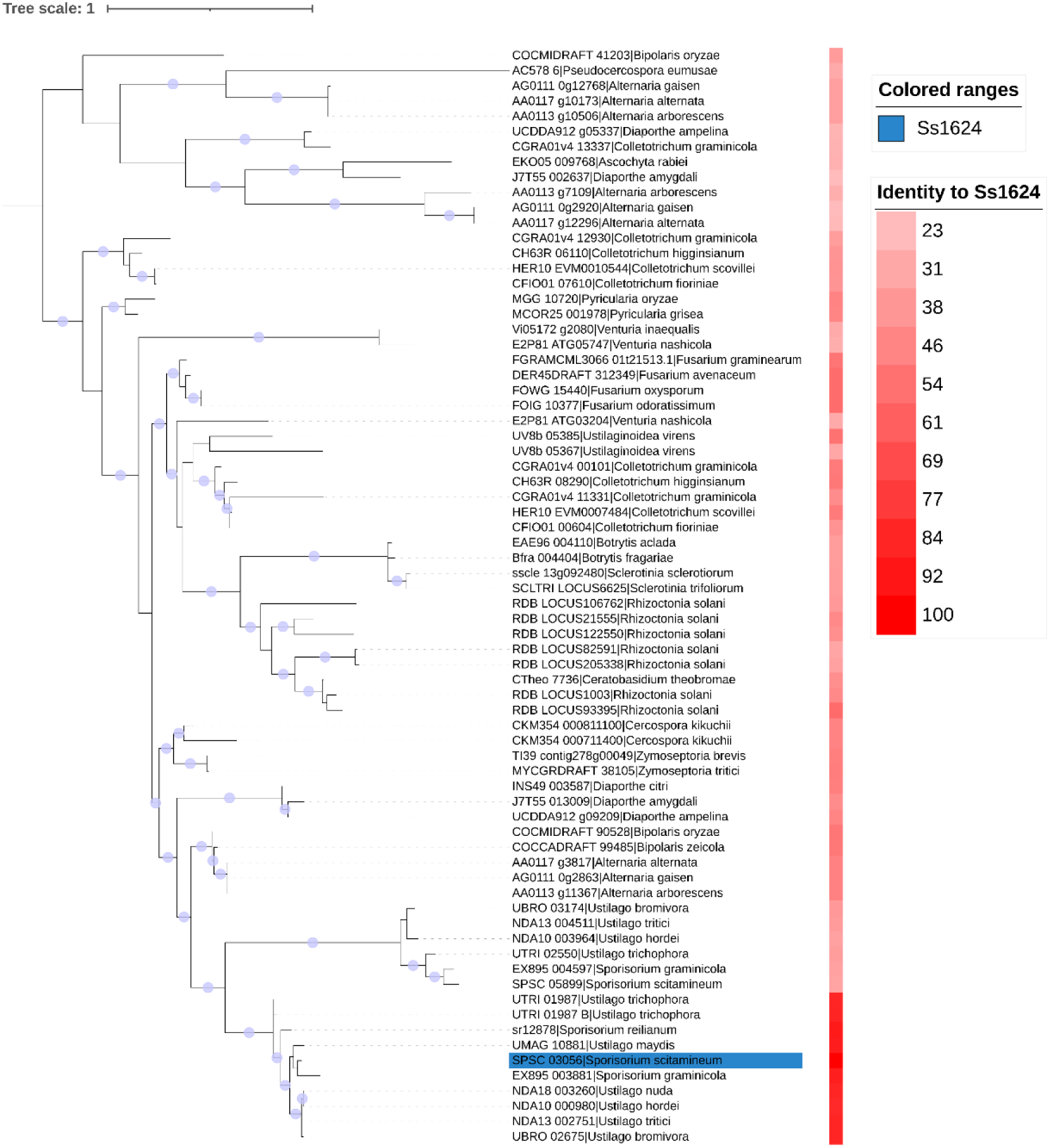
Phylogenetic analysis and evolutionary conservation of the fungal effector Ss1624. Homologs of Ss1624 (OrthoDB v12.2 orthogroup ‘400260at4751’) were retrieved and analyzed. Maximum-likelihood phylogeny (IQ-TREE 3) revealed that Ss1624 is widely distributed across fungal pathogens, exhibiting exceptional sequence conservation particularly within the *Ustilago* and *Sporisorium* genera.

**Supplemental Table 1.**
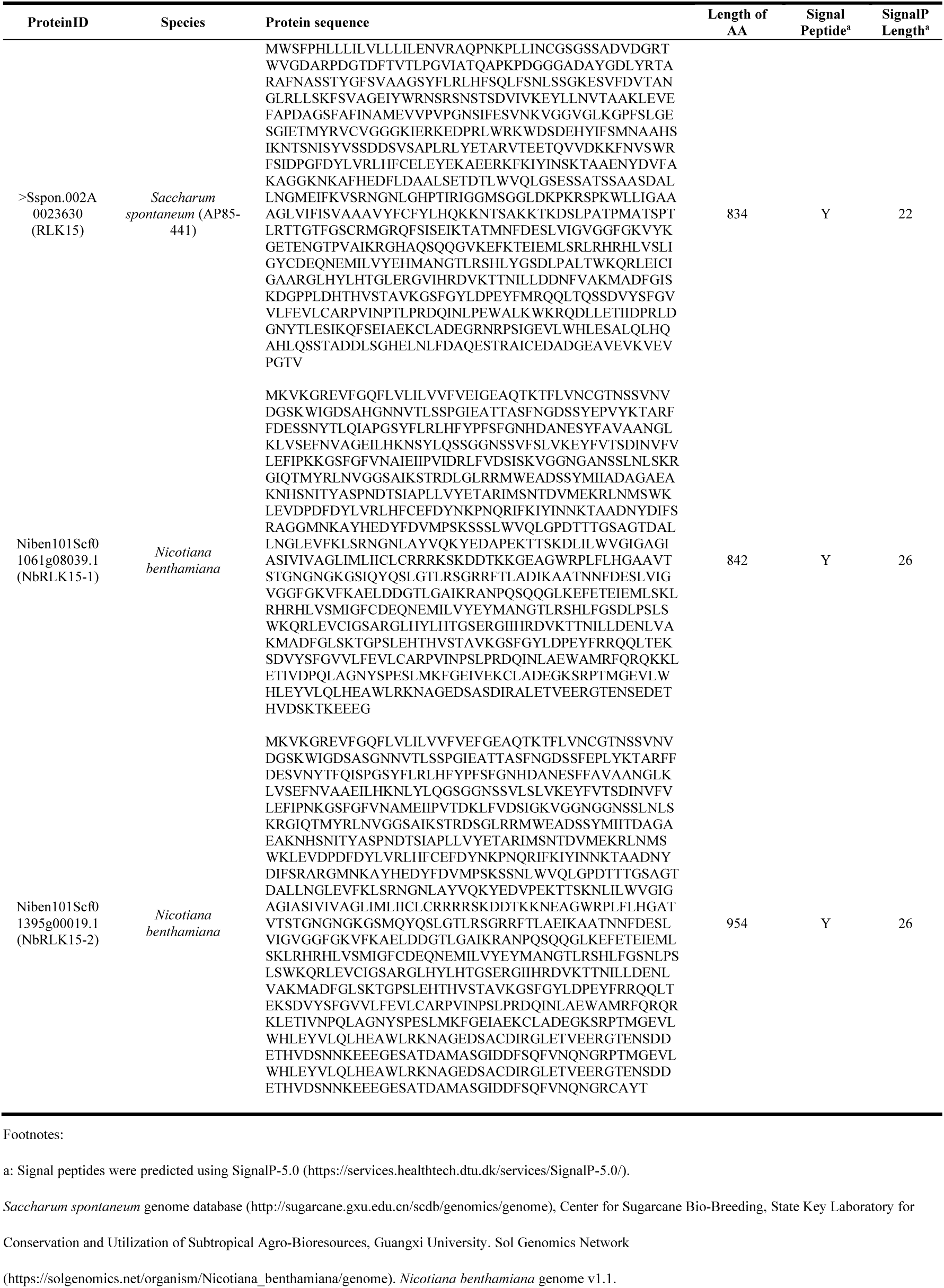
Homologous proteins of RLK15 in *N. benthamiana*.

**Supplemental Table 2.**
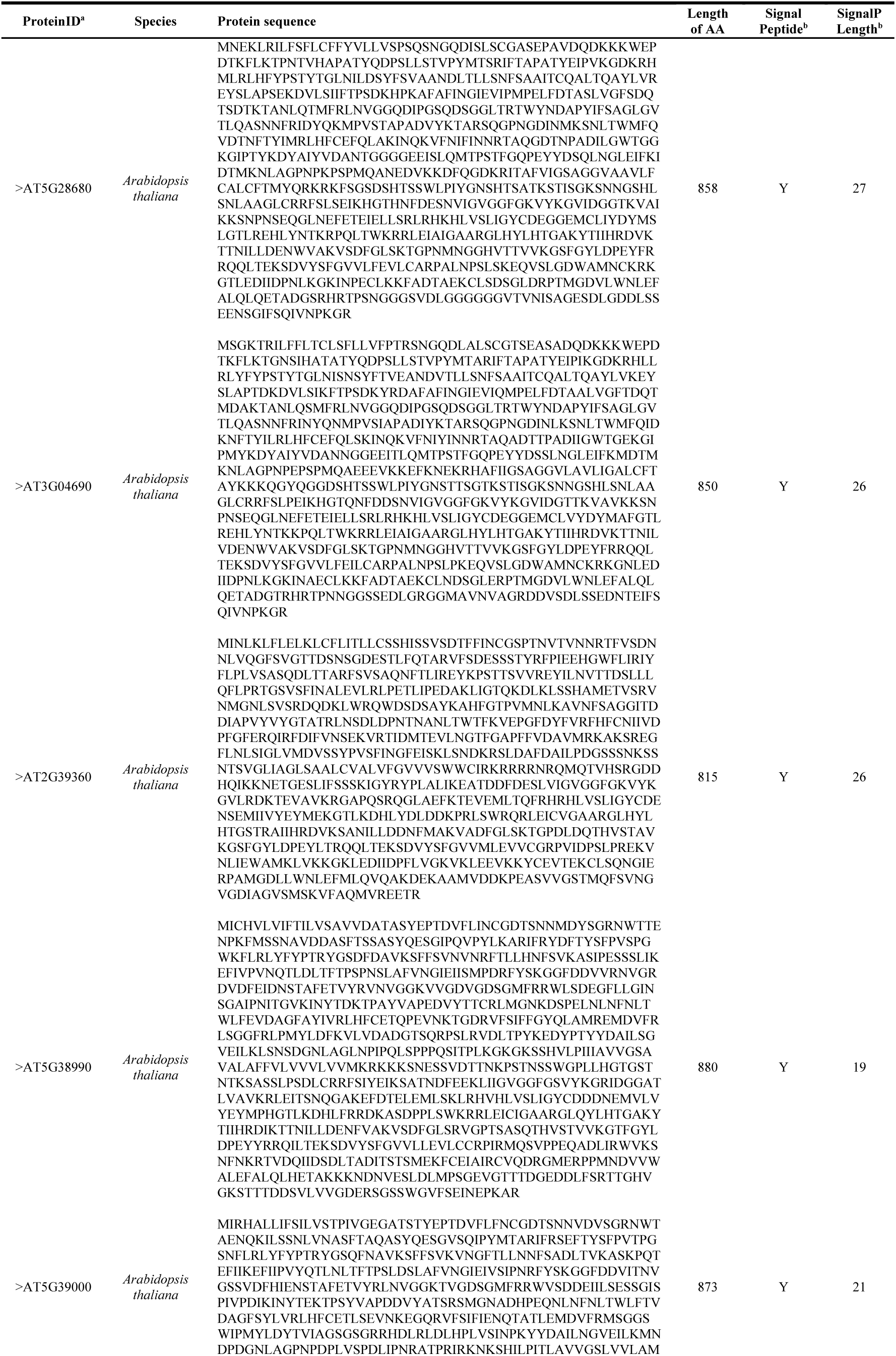
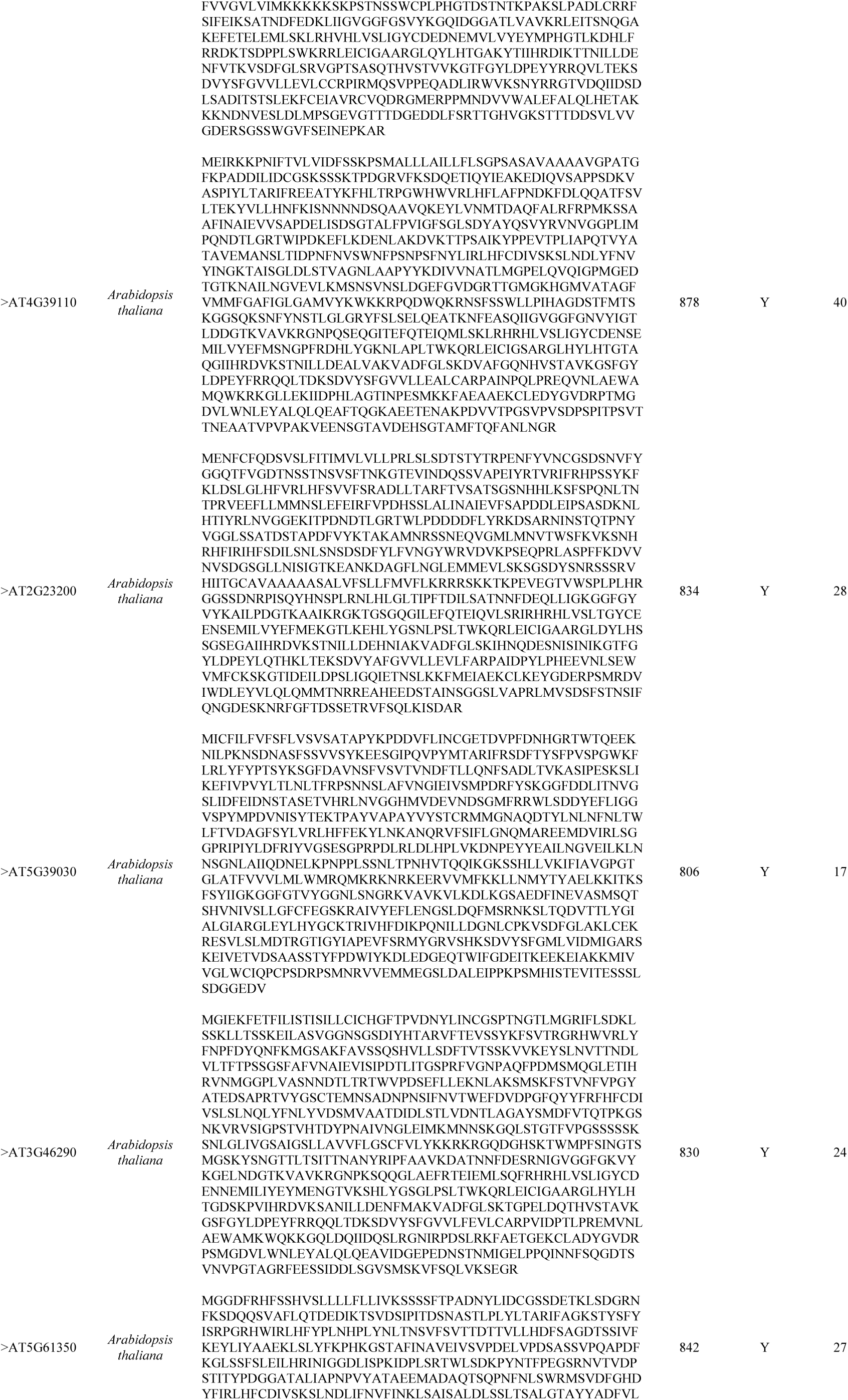
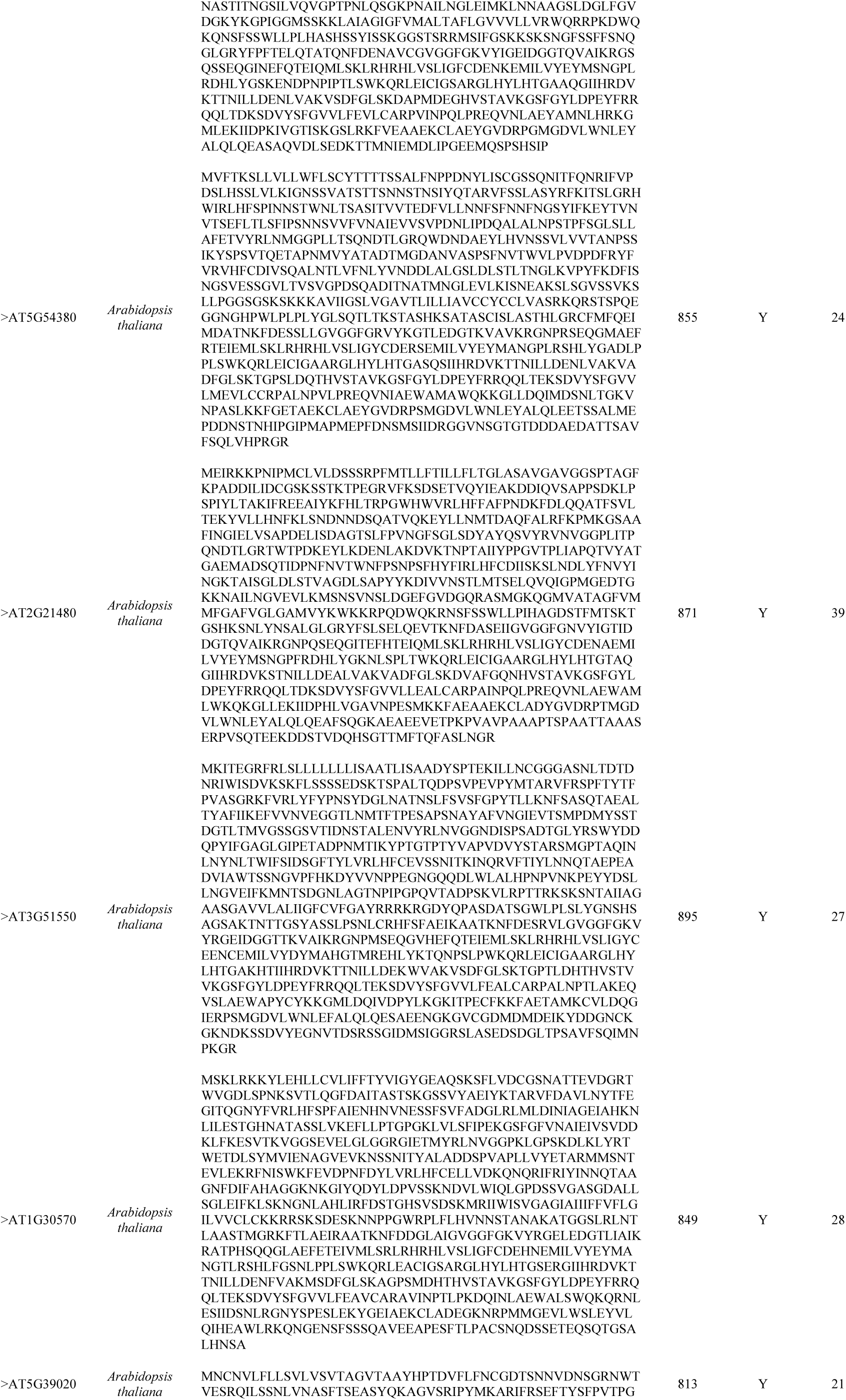
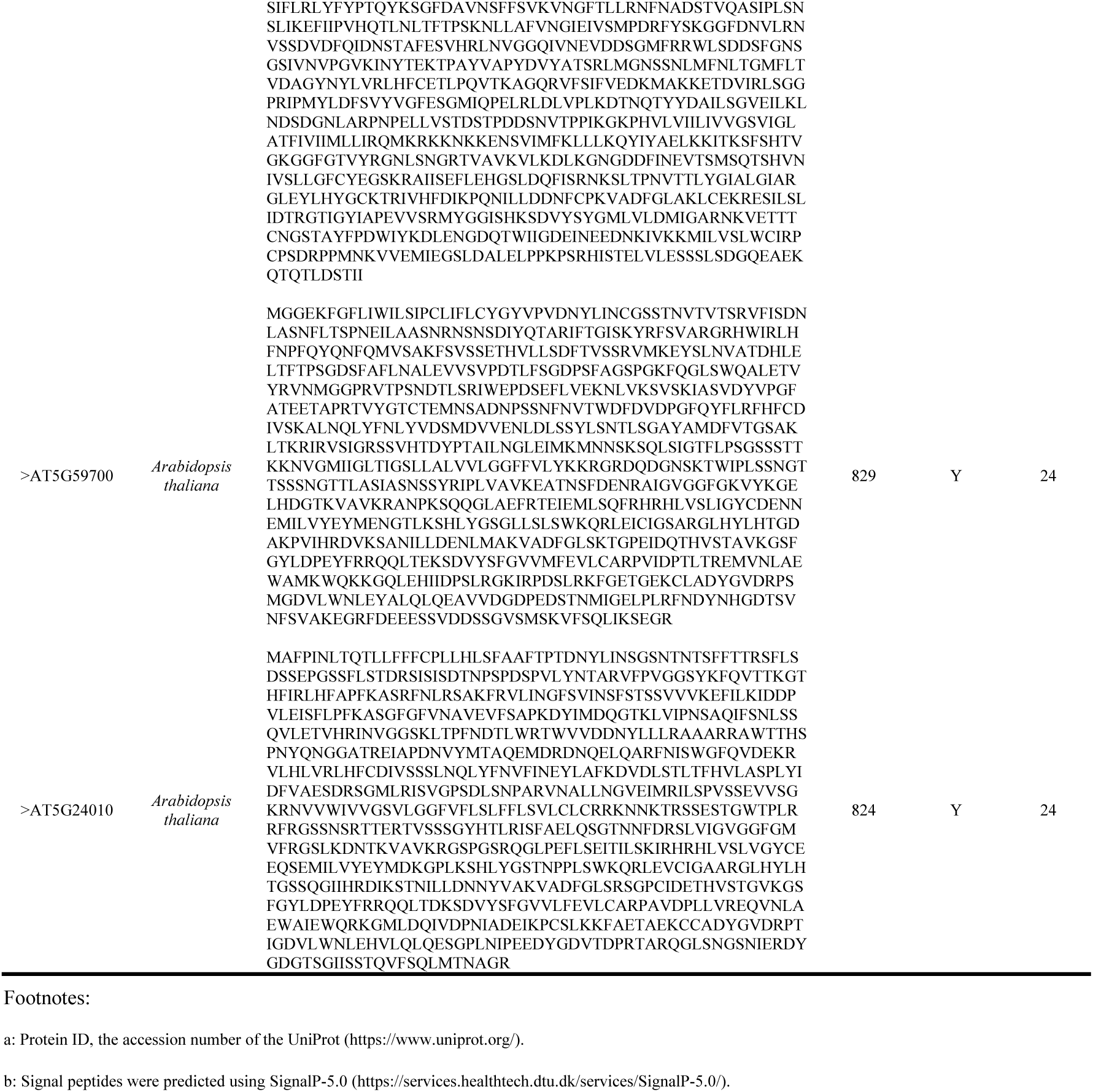
Members of the CrRLK1L Family in *Arabidopsis thaliana*.

**Supplemental Table 3.**
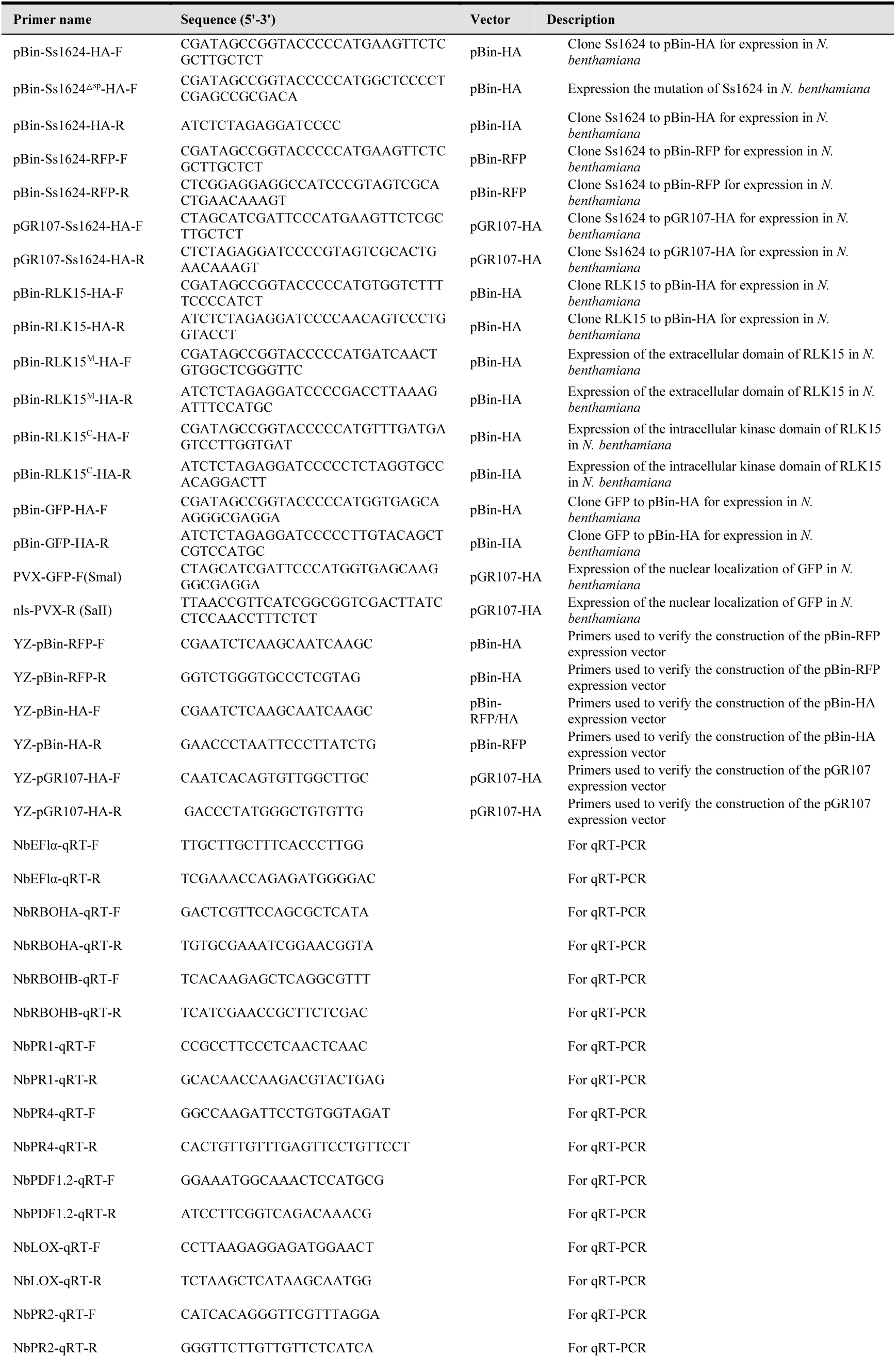
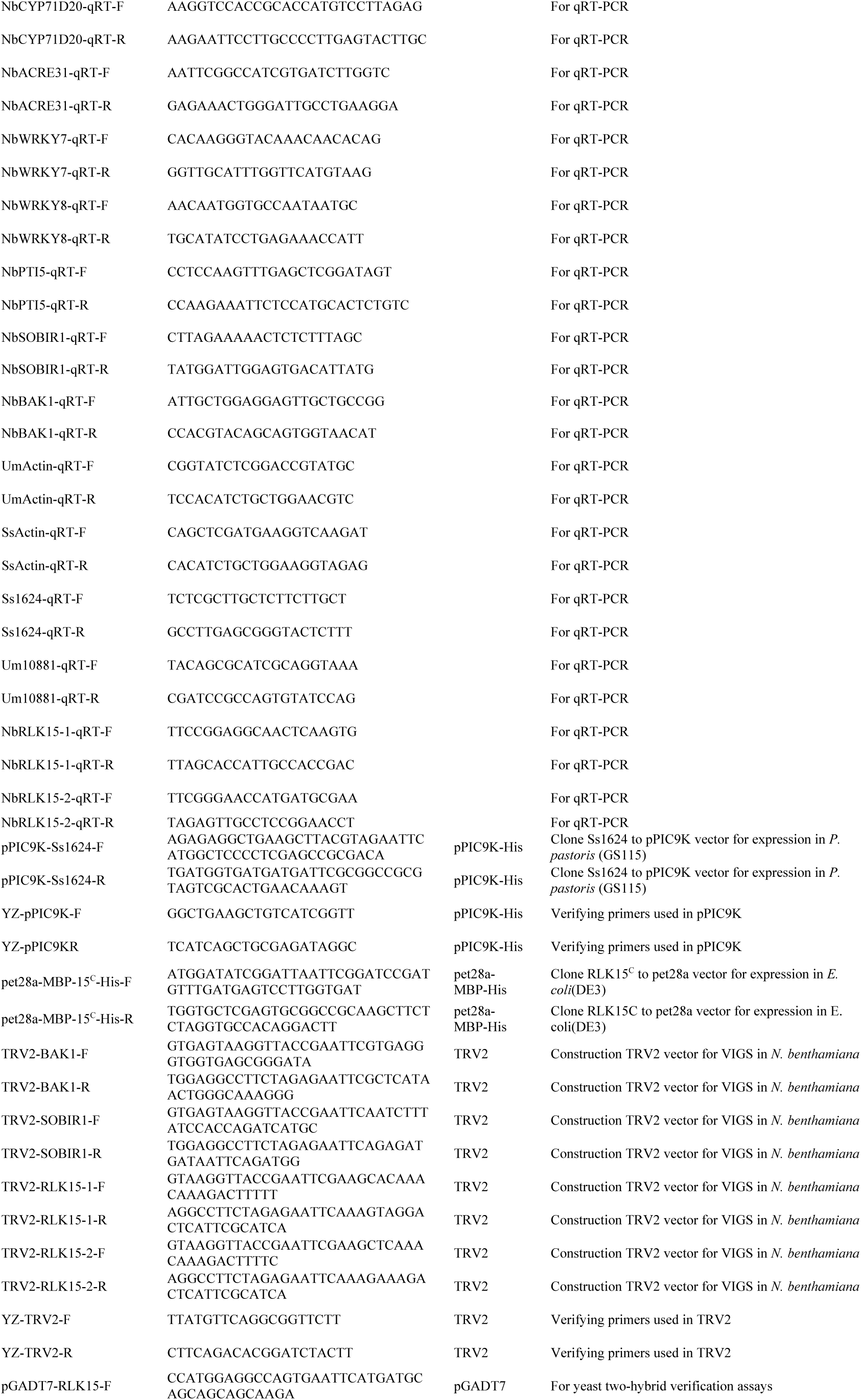
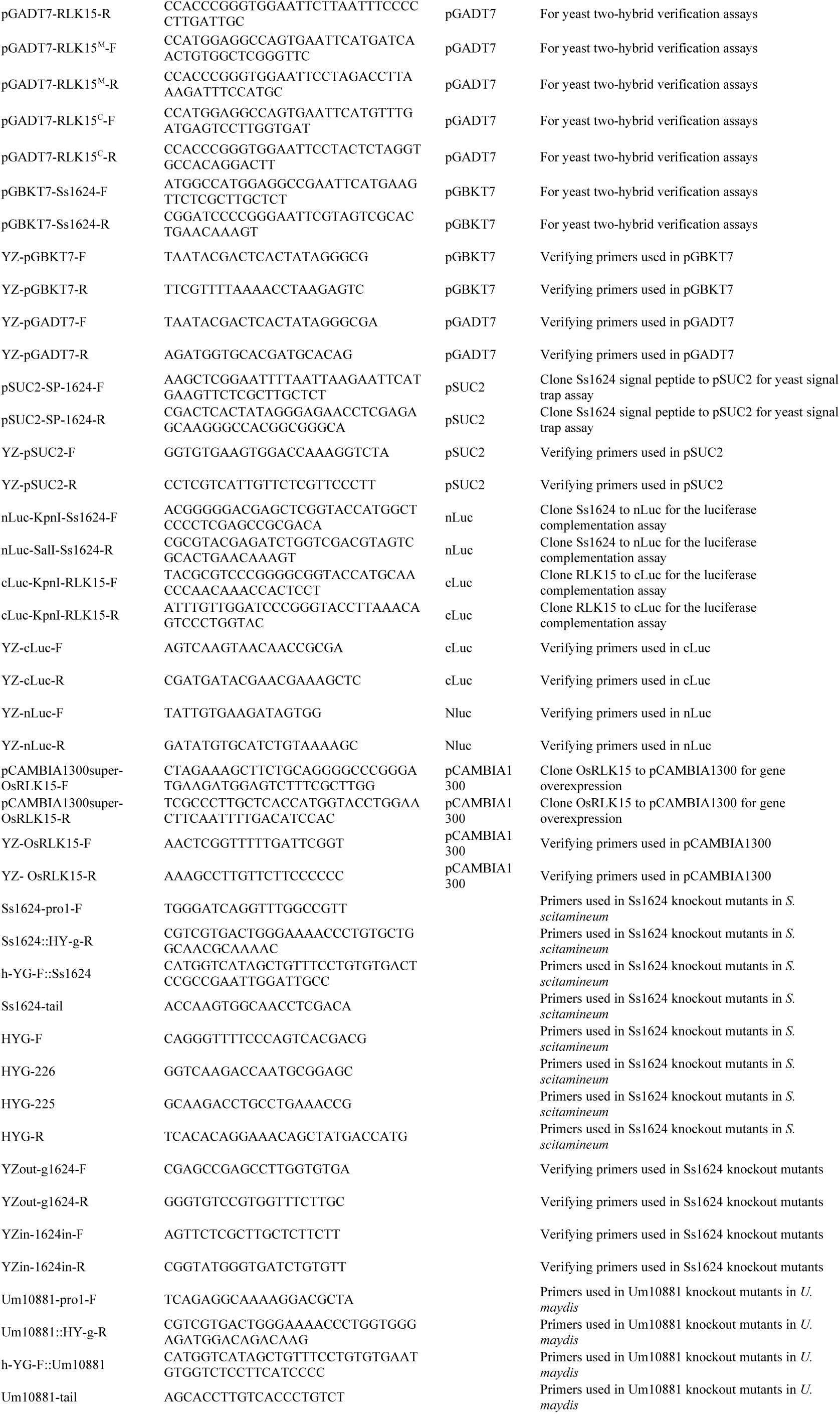
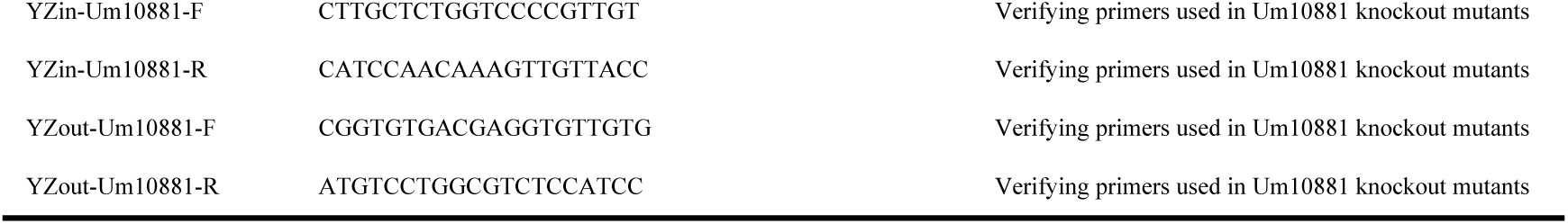
The primers used in this study.

**Supplemental Table 4.**
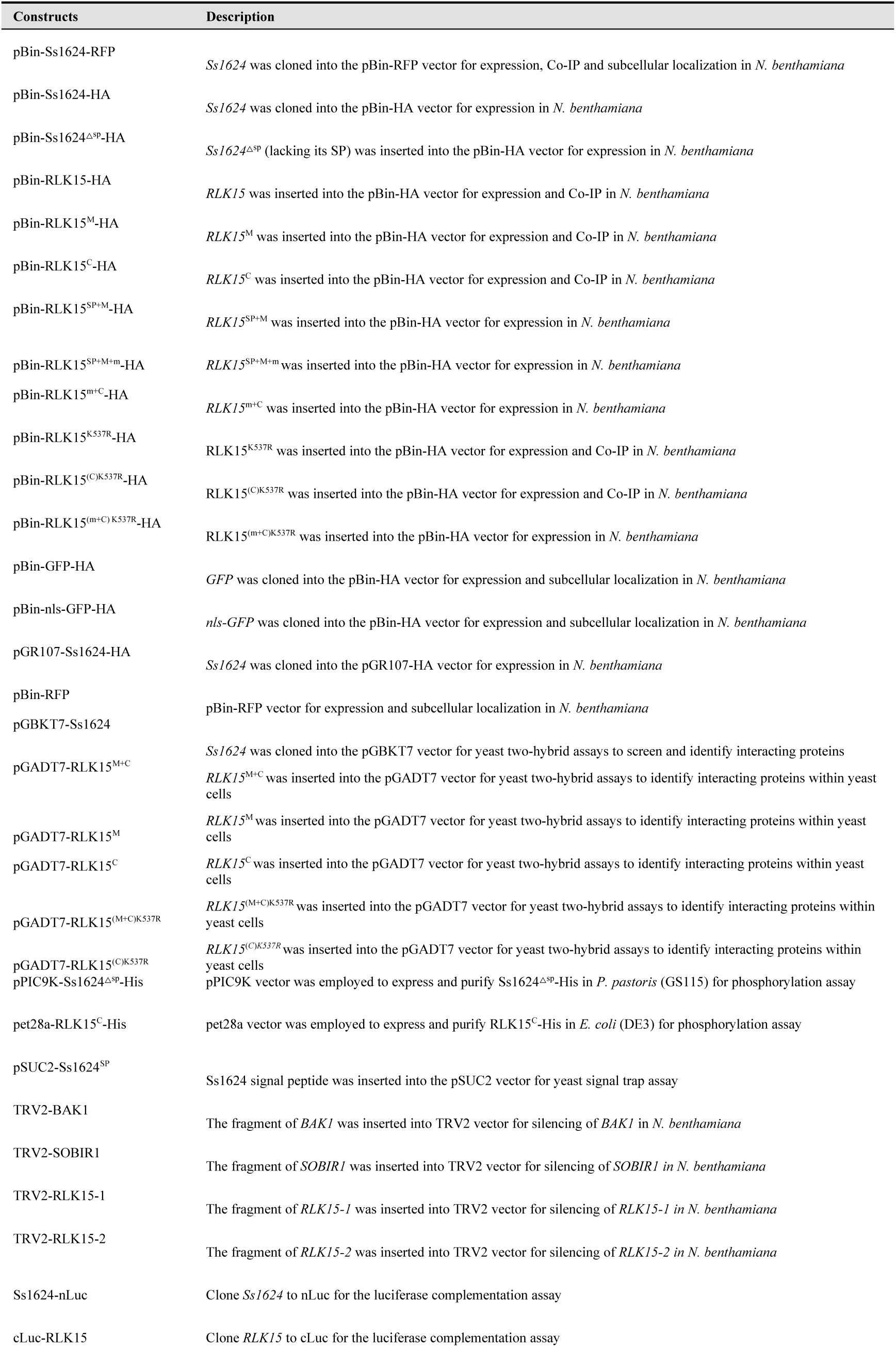
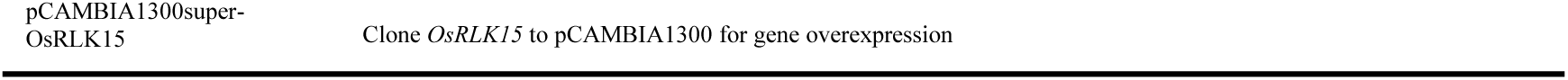
Plasmid constructs used in this study.

## Funding

This work was supported by the National Key Research and Development Program (No. 2024YFD1200800), the National Natural Science Foundation of China (No. 32472498 and U23A20148).

## Author Contributions

C.C. and L.L. designed research; L.L., Y.L., H.J., L.Z., S.T., M.Y., E.C. performed research; Z.J., L.Z and Y.D. contributed new reagents/analytic tools; L.L., Y.L., and X.Z. analyzed data; and L.L., C.C., X.Z., Y.G., and G.S. wrote the paper.

## Acknowledgements

We thank Prof. Rongxing Shen, Prof. Zeling Xu and Prof. Lisheng Liao from South China Agricultural University for their critical reading and valuable discussions of this manuscript. We thank Prof. Guanghui Kong and Gan Sha (South China Agricultural University, China) for kindly offering plant materials and plasmids.

## Declaration of Interests

The authors declare no competing interest.

## Supporting Information

## Notes

### Competing Interest Statement

The authors have declared no competing interest.

