## Supplementary figures and images for "Receptor-like kinase RLK15 binds a fungal RNase effector to enhance autophosphorylation and activate host defense"

### supplementary data 1

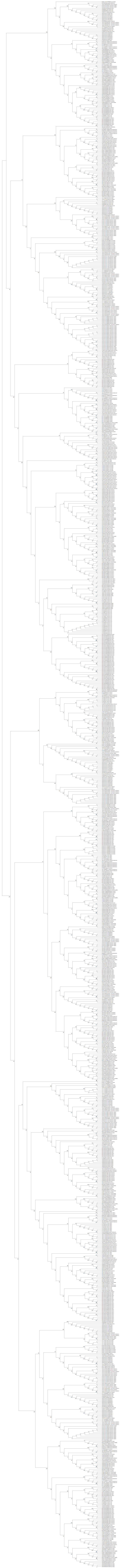
